# Point mutations and complex variants impact gene expression and addiction-related behaviors in Heterogeneous Stock rats

**DOI:** 10.64898/2026.09.10.750683

**Authors:** Denghui Chen, Katarina A. Cohen, Khai-Minh H. Nguyen, Helyaneh Ziaei Jam, Ryan J. Eveloff, Yizhi Wang, James Guevara, Thiago Missfeldt Sanches, Milad Mortazavi, Daniel Munro, Oksana Polesskaya, Jonathan L. Sebat, Melissa Gymrek, Abraham A. Palmer

## Abstract

While various variants, including single nucleotide polymorphisms (**SNPs**), small insertions/deletions, short tandem repeats and structural variants, drive individual genetic differences, their influence on gene expression and complex traits remains unclear. We used short- and long-read sequencing to generate a comprehensive variant catalog in Heterogeneous Stock (**HS**) rats and performed joint cis-expression quantitative trait loci (**cis-eQTL**) mapping across five brain regions. We found that non-SNP variants accounted for over 50% of lead regulatory associations, many of which were poorly tagged by nearby SNPs using linkage disequilibrium (**LD**). Comparison between joint and SNP-only analyses showed that over 46% of shared eQTL genes had a non-SNP lead cis-eQTL, and fewer than half were in strong LD with the corresponding lead eSNP. Linking joint cis-eQTLs to complex trait associations identified mechanisms missed by SNP-only approaches, highlighting the importance of incorporating diverse variant types into genetic studies and providing a foundational resource for the HS rat community.

## Introduction

Understanding how genetic variation shapes complex traits is a central challenge in genetics. While studies have focused on single nucleotide polymorphisms (**SNPs**), individual genomes are also defined by other types of variants, such as small insertions/deletions (**indels**), short tandem repeats (**STRs**), and structural variants (**SVs**)^1–3^. Because SNPs are abundant and technically easier to genotype, they are often used as proxies to “tag” other variant classes through linkage disequilibrium (**LD**). However, this tagging is frequently incomplete, particularly for STRs and SVs, which often possess higher mutation rates or greater structural complexity than SNPs^1,4,5^. Consequently, while genome-wide association studies (**GWAS**) have uncovered thousands of loci, pinpointing causal variants and mechanisms remains difficult^6–10^. Expression quantitative trait locus (**eQTL**) mapping analysis identifies variants significantly associated with expression of genes (**eVariants**), and provides a potential mechanism by which a variant alters gene expression, thereby influencing more complex traits that are the focus of GWAS^8,11–15^. Yet, by focusing exclusively on SNPs, many GWAS and eQTL studies overlook the regulatory contributions of non-SNP variants. Growing evidence indicates that other types of variation also play important roles in gene regulation and complex traits ^1,2,5,16–21^. Incorporating these diverse variant types is essential for uncovering the biological mechanisms missed by SNP-only approaches and for building a more complete view of the genetic architecture of complex traits.

The N/NIH Heterogeneous Stock (**HS**) rat provides a powerful model for investigating the genetic basis of behavioral and physiological traits^22–34^. Derived from the outcrossing of eight inbred founder strains (ACI/N, BN/SsN, BUF/N, F344/N, M520/N, MR/N, WKY/N, and WN/N), HS rats harbor fine-grained recombination and rich allelic diversity, enabling high-resolution mapping of complex traits^35–39^. The outbred HS rats have been maintained for over 100 generations, and have been widely used to study a range of phenotypes, including addiction-related behaviors^40–46^. Previous efforts in HS rats have focused on SNPs, leaving the contribution of other variant types unexplored^47–49^. Advances in sequencing technologies now enable comprehensive discovery and genotyping of diverse variant types^3,50–54^, offering an opportunity to revisit gene regulatory architecture and reveal new genetic mechanisms behind complex traits.

Here, we used previously generated Illumina whole-genome sequencing (**WGS**) short reads and more recently generated Pacific Biosciences (**PacBio**) high fidelity (**HiFi**) long reads on the eight HS rat founder strains and eighty-six outbred HS rats to build a comprehensive multi-type variant catalog for the HS rat population. Leveraging this resource, we extended our previous eQTL mapping study on only SNPs^48^ to diverse variant types by performing joint local eQTL (**cis-eQTL**) mapping analyses across five brain regions relevant to addiction. By directly comparing these results to a traditional SNP-only approach, we quantified the regulatory insights gained by incorporating diverse variant classes. We then utilized LD to link our joint eQTL results to published GWAS signals for addiction-related behavioral traits, uncovering previously unexplored molecular mechanisms and candidate genes. Finally, our multi-type variant and eQTL catalogs are publicly available as a foundational genetic resource for the HS rat community and provide a framework for incorporating complex variation into functional genomics. (Figure 1).

**Figure 1.**
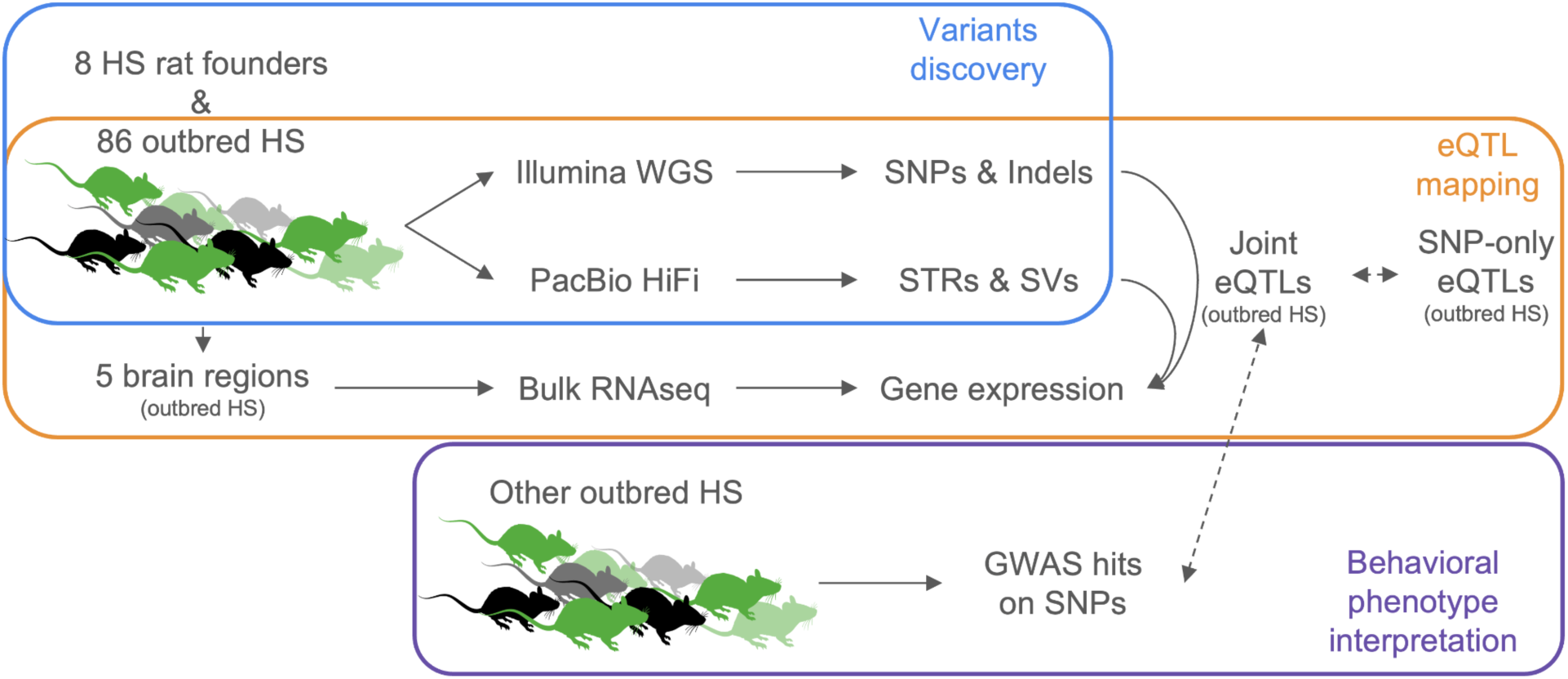
Overview of study design. We generated a comprehensive catalog of genetic variants in the HS rat population, performed eQTL analyses and investigated potential links between eVariants and GWAS signals in addiction-related behavioral traits.

## Results

### Creating a catalog of SNPs, indels, STRs and SVs for HS rats

Using Illumina WGS short reads to detect SNPs and indels (see Methods, Figure S1 and S2), and PacBio HiFi long reads to detect STRs and SVs (see Methods, Figure S1, S3 and S4), we generated a comprehensive catalog of genetic variants for the HS rat population based on the eight HS inbred founders and 86 treatment naive outbred HS rats^48^. In total, we identified 8,984,328 SNPs, 1,691,412 indels, 237,021 STRs and 96,630 SVs in the HS rat population using rat genome mRatBN7.2. These polymorphic variants span multiple subtypes, including 824,790 indel insertions (**INDEL INSs**), 866,622 indel deletions (**INDEL DELs**), 19,561 short tandem repeats with 1 bp length repeat unit (**STR homopolymers**), 217,460 short tandem repeats with repeat unit length of 2-6 bp (**STR non-homopolymers**), 52,921 SV insertions (**SV INSs**), 43,388 SV deletions (**SV DELs**), 151 SV inversions (**SV INVs**), and 170 SV duplications (**SV DUPs**) (Figure 2A).

**Figure 2.**
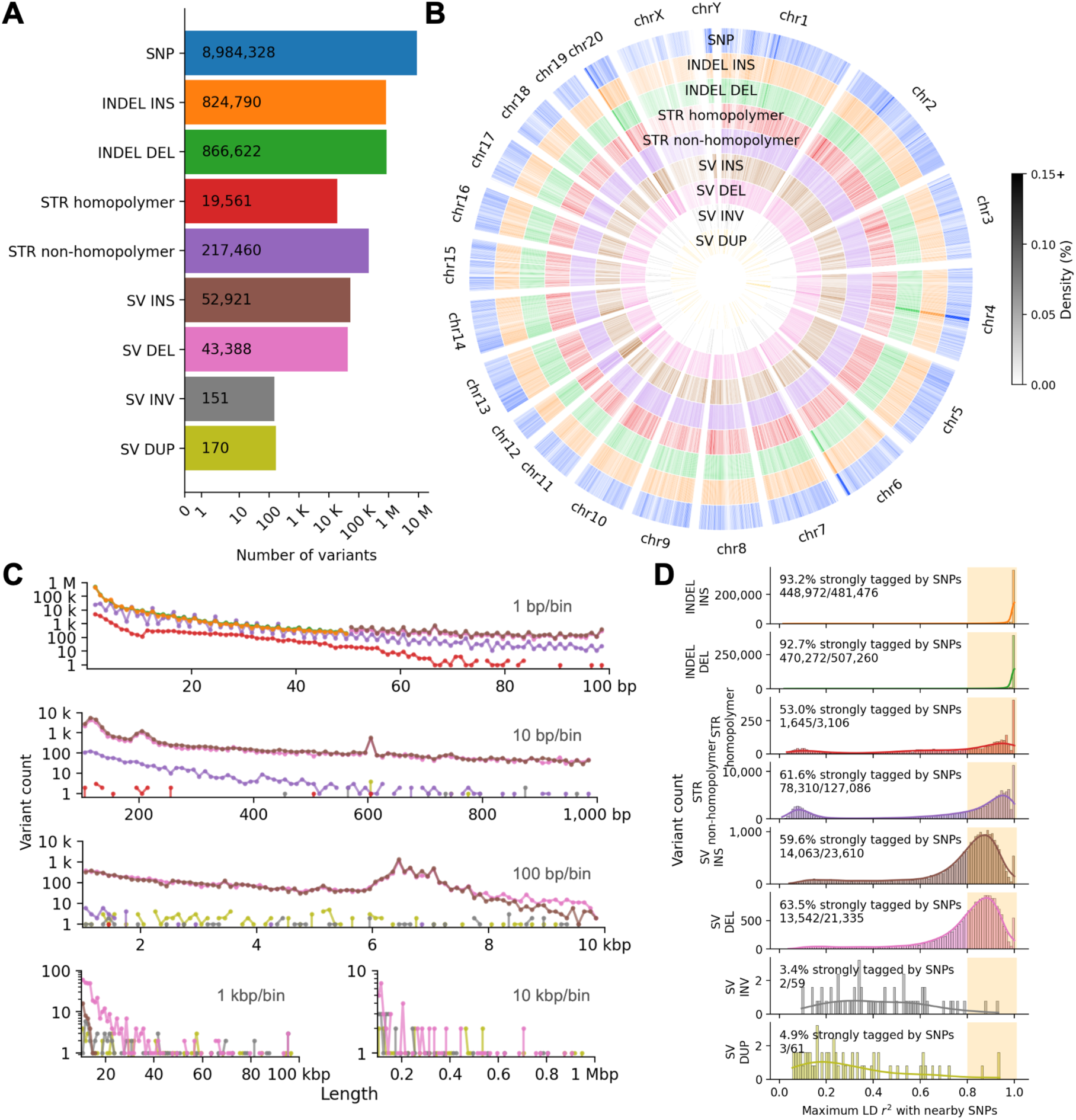
Variant discovery in HS rat population. **A.** Number of variants discovered for each variant class when compared with the Rattus norvegicus mRatBN7.2 reference genome. The X-axis is on a log scale. **B.** Circus plot showing variant density distribution on each chromosome with one-megabase windows. **C.** Length distribution of non-SNP variants. STR length represents the maximum number of base-pair differences from the reference at each locus. Same color legend as **A**. The X-axis is on a log scale. **D.** Maximum linkage disequilibrium between an autosomal non-SNP variant and any SNP within one megabase. Highlighted areas indicate variants in strong LD with nearby SNPs (r^2^ ≥ 0.8).

Using orthogonal sequencing modalities (see Methods, Figure S5 and S6), we evaluated STR and SV variant calls in the 8 founders and the 86 outbred rats. The PacBio HiFi STR call set in our catalog showed genotype concordance of 96.93% for the 8 founders and 92.54% for the 86 outbred rats when compared to the Illumina short-read WGS validation set (Figure S5). For SVs, the PacBio HiFi call set achieved 87.68% genotype concordance against a validation set generated from PacBio HiFi and Oxford Nanopore long reads for a single outbred HS rat (Figure S6).

The different types of variants were relatively uniformly distributed throughout the genome, as shown by the Circos density plot (Figure 2B). However, several regions, particularly on chromosomes 2, 4, 6, and 20, exhibited locally elevated variant densities across multiple variant types (Figure 2B and S7). These peaks may reflect regions of high genomic instability or structural complexity^55–61^.

We next assessed the length distribution of indels, STRs, and SVs (Figure 2C). We restricted indels to those with a length shorter than 50 bp, whereas SVs ranged from 50 bp to 1 Mb. We also excluded indels and SVs with ≥ 50% reciprocal overlap with STRs to ensure mutually exclusive variant sets. All variant types showed a decline in abundance with increasing length and were plotted on a log scale (Figure 2C). We observed peaks for SVs around 200 bp, 600 bp, and 6.5 kb. These features likely correspond to common repetitive elements in the rat genome, such as short interspersed nuclear elements (SINE), truncated endogenous retroviral insertions, and long interspersed nuclear elements (LINE), respectively^61–66^. In terms of total sequence affected, indels accounted for 5,482,480 bp of variable sequence (0.2071% of the reference genome), STRs accounted for 2,268,513 bp (0.08567%), and SVs accounted for 142,786,092 bp (5.392%). This highlights the disproportionate impact of structural variants on total genomic variation despite their lower frequency.

Given that most prior GWAS in rats rely on SNPs, we evaluated how well SNPs tag other types of variants through linkage disequilibrium. For each autosomal non-SNP variant, we computed their maximum LD r^2^ with any individual SNP within a one-megabase window (Figure 2D). SNPs were highly effective in tagging small indels, with 93.2% of insertions and 92.7% of deletions showing strong LD with one or more nearby SNPs (r^2^ ≥ 0.8). STRs were moderately well tagged, with 53.0% of homopolymer and 61.6% of non-homopolymer STRs reaching this threshold (r^2^ ≥ 0.8). SV insertions and deletions also showed moderate tagging rates, 59.6% and 63.5%, respectively. SV inversions and duplications were rarely tagged, with only 3.4% and 4.9% of variants in strong LD with SNPs. The low tagging rates of SV inversions and duplications should be interpreted cautiously due to the low number of SV inversions and duplications retained in the analysis. This may be because the SV caller classifies most 1kb-10kb duplications as insertions, and SV duplications may reside in segmental duplication regions. Nevertheless, these findings indicate that although SNPs capture many indels and some STRs and SVs, many STRs and SVs are not effectively tagged by SNPs and therefore their contributions to traits studied by SNP-based GWAS may have been underestimated or completely missed.

### Joint eQTL mapping

Leveraging the comprehensive variant catalog described above, we performed cis-eQTL mapping analyses in the 86 outbred HS rats across five addiction-relevant brain regions: prelimbic cortex (**PL**), infralimbic cortex (**IL**), orbitofrontal cortex (**OFC**), nucleus accumbens core (**NAcc**) and lateral habenula (**LHb**). In this study, we only focused on autosomal genes and variants. After quality control, we retained per tissue a mean of 19,308 genes and 6,428,120 variants across 78 samples (Table S1). On average, 98,004,145 variant-gene pairs were tested per tissue within a ±1 Mb window.

Using the approach described in Methods, we identified 4,020,084 unique cis-eVariants associated with 10,579 eGenes in at least one tissue at a 5% false discovery rate (Table S2). The quantile-quantile plot shows a clear deviation from the null expectation across all tissues, indicating a substantial number of significant variant-gene associations (Figure 3A). These associations span all tested variant types, and the distribution of eQTL signals was consistent across tissues (Figure S8). Furthermore, pairwise correlations of shared eQTLs’ effect sizes between tissues revealed strong cross-tissue similarity, especially between PL, IL and OFC (Spearman rho = 0.94-0.96), while LHb and NAcc showed slightly lower concordance (rho = 0.90-0.92) (Figure 3B). The overall high concordance likely reflects both shared regulatory architecture (genetic factors) and the fact that tissues were collected from the exact same individuals (environmental factors). The clustering pattern of cross-tissue correlations is similar to tissue similarity patterns previously observed using principal component analysis of gene expression in this cohort^48^.

**Figure 3.**
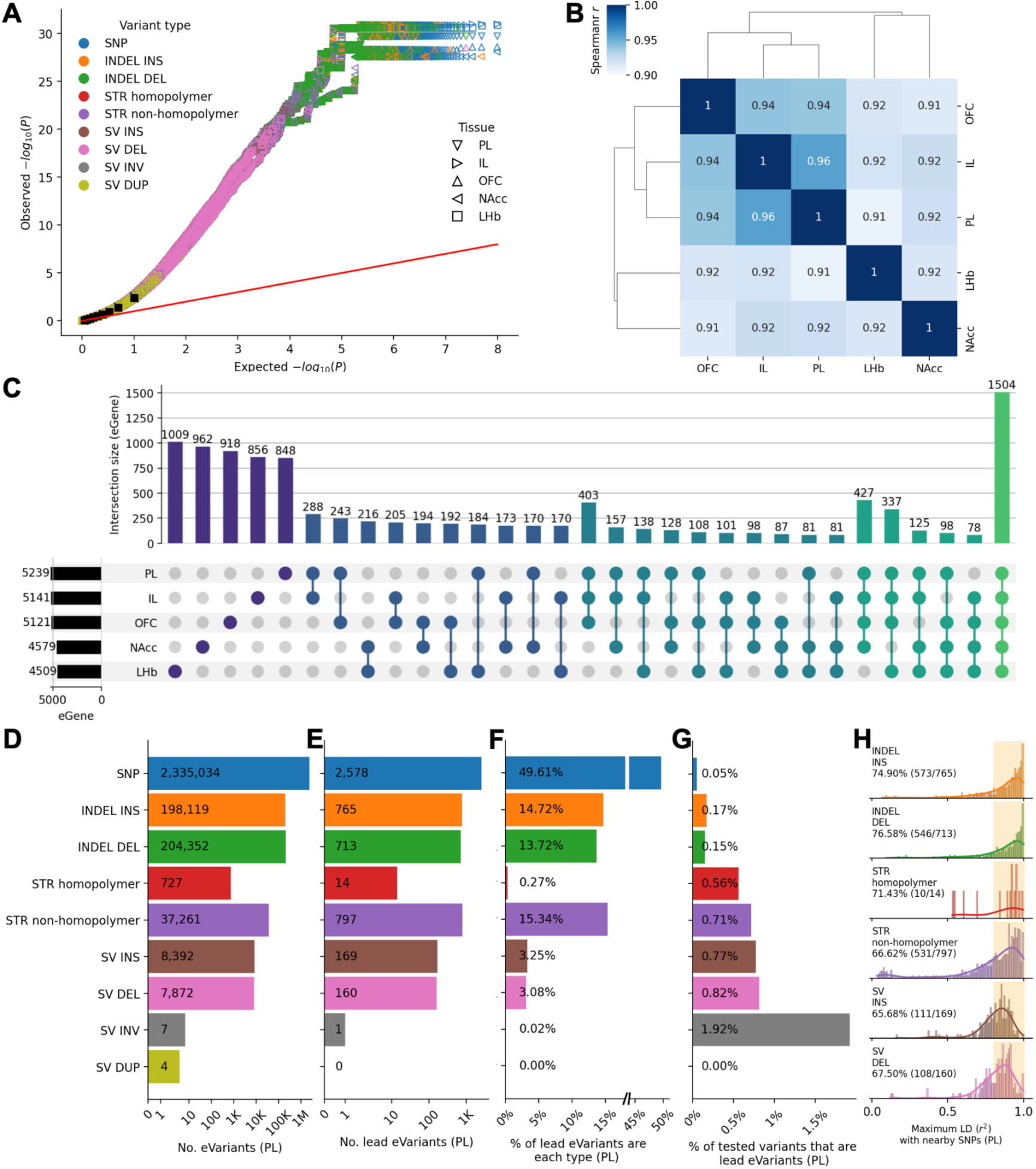
Joint eQTL analyses on five brain regions from 86 outbred HS rats. **A.** The quantile-quantile plot of observed p values for each variant by gene test compared against the expected uniform distribution for each tissue. Black squares denote deciles of the distributions. **B.** Shared eQTLs’ effect sizes correlations across tissues. **C.** Upset plot showing overlap of eGenes across tissues. **D.** Number of eVariants for PL tissue across variant types. **E.** Number of lead eVariants for PL tissue across variant types. **F.** Percent of lead eVariants that are each variant type for PL tissue. **G.** Percent of tested variants from each variant type that are lead eVariants for PL tissue. **H.** Maximum linkage disequilibrium distribution between a non-SNP lead eVariant and SNPs within one megabase for PL tissue. Highlighted areas indicate variants in strong LD (r^2^ ≥ 0.8) with nearby SNPs. Percentages indicate the proportion of lead eVariants that are in strong LD with nearby SNPs. Similar data for the 4 other tissues are shown in Figure S9.

We next examined the overlap of eGenes across tissues. Out of the 10,579 unique eGenes identified, 4,593 were specific to a single tissue, whereas 1,504 eGenes were detected in all five brain regions (Figure 3C). In line with our previous observation, PL, IL and OFC shared the largest amount of eGenes in both two and three tissues combinations, which is likely because of similar transcriptional profiles among these cortical tissues^48^. A large portion of eGenes were tissue-specific, which is also consistent with our prior findings in this cohort. The tissue-specific patterns may reflect biologically meaningful regulatory differences across brain regions, though some may also arise from statistical noise or differences in power due to higher or lower expression among the 5 tissues^48^.

We then summarized eVariants statistics by variant type. Millions of eVariants were detected in each tissue, with SNPs comprising the largest fraction, followed by indels, STRs and SVs (Figure 3D, S9A and Table S2). This distribution is likely due to the underlying abundance of each variant type in the genome, as well as differences in the number of variants tested in the joint eQTL mapping analyses. To prioritize putative causal variants, we identified the most significant eVariant for each eGene in each tissue as the lead eVariant. Across tissues, there were 187 to 248 tie cases (4.16% to 4.76% of the eQTLs) where multiple eVariants had identical *P*-values (Figure S10). In such cases, we prioritized based on functional annotation with the order of exonic, intronic, intergenic, followed by variant type with the order of SNP, indel, STR, SV. As expected, SNPs accounted for the largest number of lead eVariants, followed by indels, STRs, and SVs (Figure 3E, S9B and Table S2). While SNPs dominated in terms of overall proportion, non-SNP variants collectively accounted for more than 50% of the lead eVariants across all five tissues (Figure 3F and S9C). Although fewer indels, STRs and SVs were considered for eQTL mapping analyses, they were more likely than SNPs to be lead eVariants. (Figure 3G and S9D). This indicates that non-SNP variants contribute greatly to gene expression regulation, and have a disproportionate influence compared to SNPs.

To assess how well SNPs capture regulatory signals driven by other variant types, we quantified linkage disequilibrium between each non-SNP lead eVariant from the joint eQTL analyses and their nearby SNPs. Across five tissues, an average of 70.22% to 71.92% of the non-SNP lead eVariants were in strong LD (r^2^ ≥ 0.8) with at least one SNP within ±1 Mb (Figure 3H and S9E). Lead eSTRs and eSVs had even lower average tagging rates, ranging from 65.85% to 68.05%, presumably due to the high mutation rate of STRs and the structural complexity of SVs, both of which decouple these variants from surrounding SNPs. While many non-SNP lead eVariants’ effects may be indirectly tagged by SNPs, a large subset lack adequate SNP proxies, particularly those signals mediated by STRs and SVs. This indicates that SNP-only eQTL analysis provides incomplete coverage of regulatory architecture and is likely to miss a significant fraction of associations, which we explore in more detail below.

### Characteristics of eVariants

We next assessed whether the identified eVariants exhibited properties consistent with known cis-regulatory architectures. All significant eQTLs and lead eQTLs showed strong enrichment proximal to their associated genes’ transcription start sites (Figure S11A), reflecting the expected localization of cis-regulatory effects near gene promoters and proximal regulatory elements. Across tissues and variant types, except for a few categories with small numbers of associations, eQTLs and lead eQTLs displayed an approximately symmetrical effect size distributions that were centered around zero (Figure S11B), representing the presence of both expression-increasing and expression-decreasing regulatory alleles.

Across tissues and variant types, eVariants and lead eVariants both showed higher non-major allele frequencies (1 - major allele frequency) on average relative to all tested variants (Figure S11C), aligning with increased statistical power to detect regulatory associations for common variants. Because bi-allelic SNPs, indels and SVs were used in this study, their non-major allele frequencies were essentially minor allele frequencies, which never exceed 0.5. However, for STRs that had more than two alleles (e.g. alleles at a given STR locus might have 2, 3 or 4 copies of the repeat), non-major allele frequencies are sometimes greater than 0.5. Further examination revealed that STRs with non-major allele frequencies greater than 0.5 were driven by their larger number of alleles (Figure S12). Consistent with statistical power considerations, eSTRs and lead eSTRs were enriched at non-major allele frequencies near 0.5 and at lower allele counts (Figure S12).

Given the unique regulatory properties of tandem repeats, we further examined whether lead eSTRs exhibited enrichment for specific repeat motifs. Enrichment analysis of lead eSTRs combined across all tissues revealed significant overrepresentation of GC-rich repeat units compared to background STRs tested in the eQTL analyses (Figure S11D and Table S3). This enrichment is consistent with prior observations that GC-rich repeats are more likely to influence transcriptional regulation, potentially through effects on chromatin structure, DNA secondary structure, or transcription factor binding^67,68^.

Finally, we assessed the functional genomic annotations of lead eQTLs for each variant type, combining signals across all tissues. Compared to all variant-gene pairs tested in the eQTL analyses, lead eQTLs were enriched across all categories that we considered, including exonic, intronic, UTR, and promoter regions (Figure S11E and Table S4). However, this enrichment likely reflects the non-uniform distribution of both eVariants and genomic annotations within the cis window. Because regulatory signals and functional annotations are both densest near transcription start sites, even non-causal lead eVariants in LD with the true causal signal would show a degree of enrichment in these features due to their genomic proximity^48^.

### Joint vs SNP-only eQTL mapping

To evaluate how adding non-SNP variants affects regulatory association discovery, we compared our joint eQTL results to traditional SNP-only cis-eQTL analyses, which excluded indels, STRs and SVs while using the same mapping pipeline. The SNP-only analyses identified 3,521,120 unique cis-eSNPs associated with 10,486 eGenes in at least one tissue at a 5% false discovery rate (Table S5). Per tissue, between 3,988 to 4,716 eGenes were identified by both the joint and SNP-only eQTL analyses, accounting for 87.09% to 90.02% of the eGenes detected in the joint eQTL analyses (Figure 4A and S13A). On average, 10.34% of the eGenes identified in the SNP-only eQTL analyses were not detected in the joint analyses. These eGenes were likely due to stricter gene-level thresholds in the joint analyses arising from larger variant sets per gene, causing borderline eGenes in the SNP-only analyses to fall below significance. Conversely, 11.03% of eGenes were uniquely identified in the joint eQTL analyses, indicating novel discoveries enabled by including indels, STRs and SVs that capture regulatory variation missed by SNPs alone.

**Figure 4.**
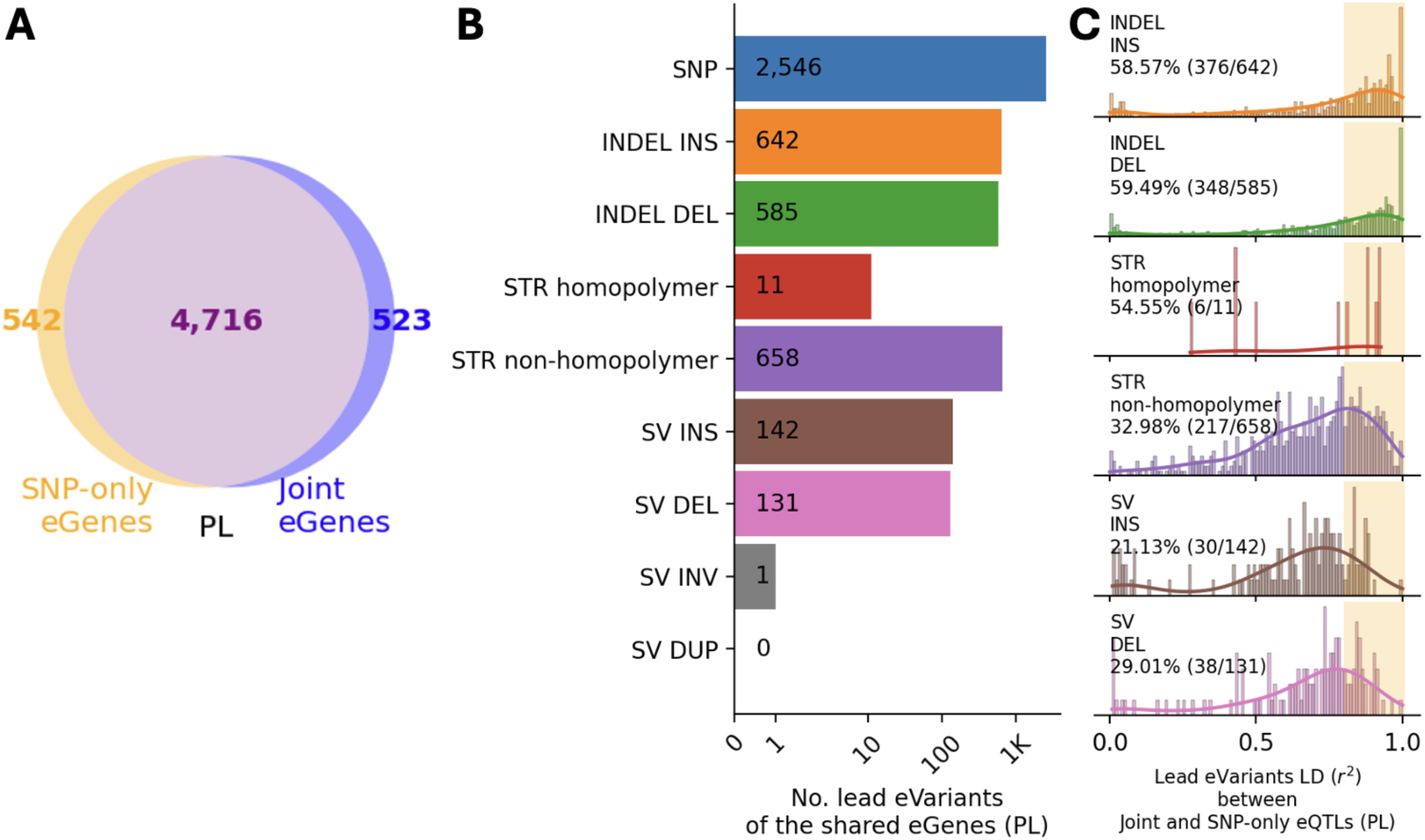
Joint versus SNP-only eQTL analyses in the 86 outbred HS rats. **A.** Venn diagram showing the intersection between eGenes detected in the joint and the SNP-only eQTL analyses for PL tissue. **B.** Number of lead eVariants of the shared eGenes across variant types in the joint eQTL analysis for PL tissue. **C.** Linkage disequilibrium distribution between shared eGenes’ lead eVariants in the joint eQTL analysis and corresponding lead eSNPs in the SNP-only eQTL analysis for PL tissue. Highlighted areas indicate variants in strong LD with r^2^ ≥ 0.8. Percentages indicate the proportion of lead eVariants in the joint eQTL analysis that are in strong LD with lead eSNPs in the SNP-only eQTL analysis. Similar data for the 4 other tissues are shown in Figure S13.

To determine whether the shared eGenes were driven by the same underlying variant, we measured linkage disequilibrium between the lead eVariants identified in the joint analyses and their corresponding lead eSNPs from the SNP-only analyses. Among the 3,988 to 4,716 shared eGenes across five tissues, 2,097 to 2,546 (52.19% to 53.99%) were mapped to a lead SNP variant in the joint eQTL analyses (Figure 4B and S13B). The remaining 46.01% to 47.81% mapped to a non-SNP lead eVariant, indicating that, when allowed to compete with SNPs, non-SNP variants frequently showed the strongest associations. We then assessed LD between non-SNP lead eVariants and their corresponding lead eSNPs. Across tissues, only 44.39% to 46.80% were in strong LD (r^2^ ≥ 0.8) (Fig 4C and S13C). STR and SV lead eVariants showed notably lower levels of LD with their corresponding lead eSNPs, ranging from 30.14% to 35.46%. This suggests that SNP-only mapping frequently prioritizes lead SNPs that do not capture the strongest association signal observed in the joint analyses.

### Biological insights gained from joint eQTL mapping

Incorporating indels, STRs, and SVs into a joint eQTL mapping framework enables the discovery of regulatory mechanisms that are difficult to capture with SNPs alone, particularly when repeats or structural changes alter transcript processing. A clear example is a 220 bp eSV deletion associated with decreased *Crybb1* expression in LHb tissue (Figure 5A, S14A). This SV overlaps exon 5 of *Crybb1*, removing 51 bp of exonic sequence and 169 bp of adjacent intron (Figure 5B). Heterozygote carriers (n=11) and a single rat that was homozygous (n=1) for the deletion showed dosage-dependent exon 5 skipping with a prominent exon 4-6 splice junction and increased RNA-seq read coverage across the intervening intronic region (Figure 5C). Because *Crybb1* has previously been implicated in human cataracts^69–71^, we next tested whether this eSV deletion is associated with *Crybb1* expression in an independent eye tissue dataset from the RatGTEx database. As expected, *Crybb1* expression was significantly higher in eye tissue than LHb (Figure S14A and S14B), providing more precise splicing quantification in eye tissue relative to LHb. Examination of the eight HS rat founders’ SV callset indicated that the WN/N inbred strain is the only carrier of this deletion. Using SNPs within the *Crybb1* gene body to identify the WN/N haplotype, we imputed the deletion into the eye RNA-seq samples (0/0 n=46, 0/1 n=5, 1/1 n=2) and observed a similar SV dosage-dependent exon 5 skipping pattern in eye tissue (Figure 5D). Lastly, we quantified percent spliced-in (**PSI**) for exon 5 in both LHb and eye tissues, and found that samples lacking the SV showed near-complete exon 5 inclusion, heterozygous carriers showed intermediate inclusion (mean PSI 77.96% in LHb; mean PSI 65.66% in eye), and homozygous carriers showed near-complete exon 5 exclusion (Figure S14C and S14D). Notably, PSI estimates in LHb should be interpreted cautiously given the very low expression level on *Crybb1*. Nonetheless, the concordant junction-level evidence and the consistent direction of effect across tissues (LHb and eye) suggest that the SV deletion disrupts splicing of exon 5.

**Figure 5.**
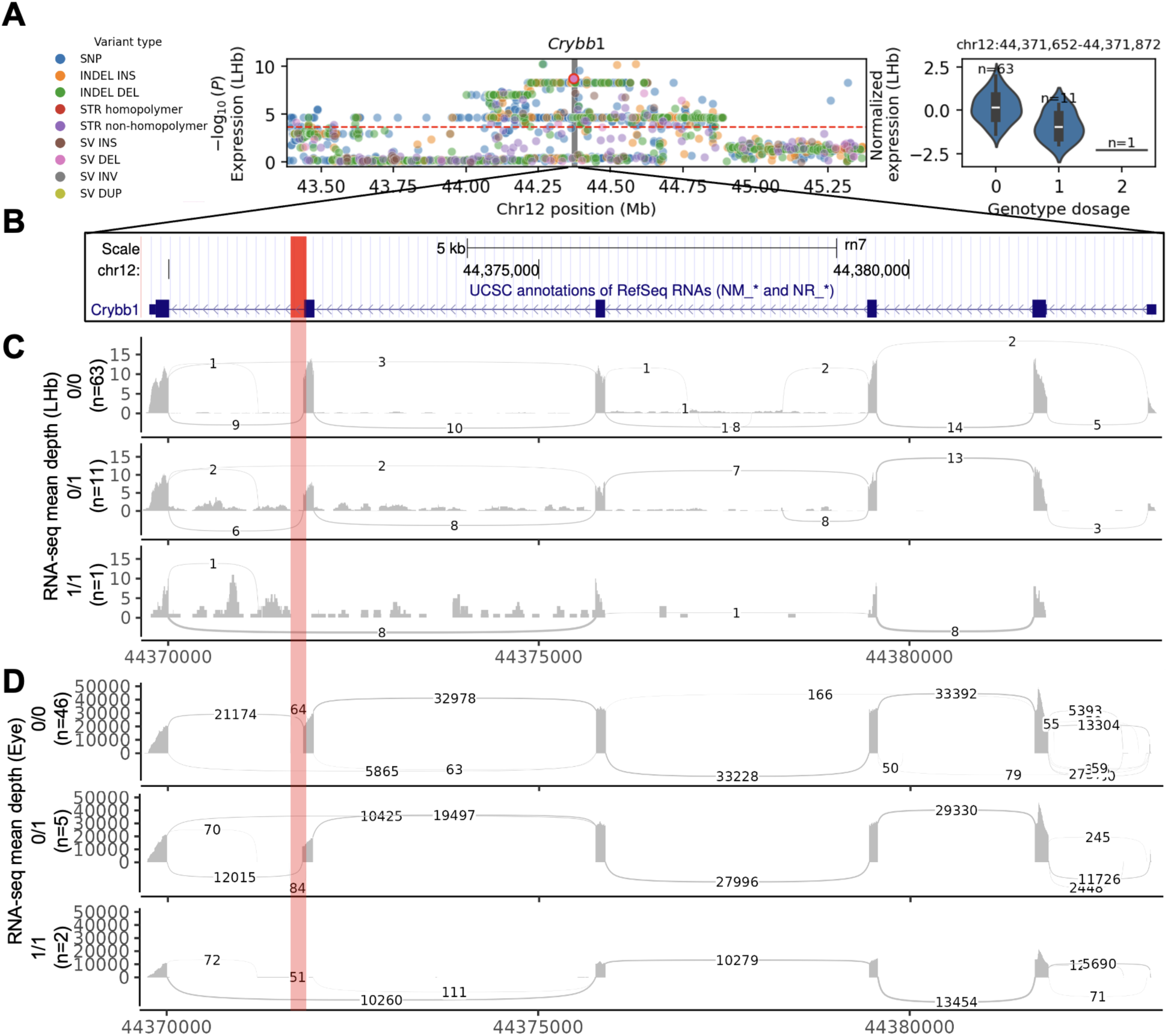
Exon-disrupting eSV affecting expression of *Crybb1*. **A.** eSV association for *Crybb1*. The left panel shows nominal -log10(*P*-values) for all tested variants in *Crybb1* in LHb tissue. The gray background indicates the genomic position of *Crybb1*. The highlighted dot denotes the exon-disrupting eSV. The right panel shows the effect plot for the SV-eQTL with genotypes on the x-axis and normalized expression on the y-axis. **B.** Genome view of the eSV locus in *Crybb1*. The highlighted band shows the genomic location of the 220 bp eSV. **C.** Sashimi plots showing that the eSV disrupts expression of exon 5 of *Crybb1* in LHb tissue. **D.** Sashimi plots showing that the eSV disrupts expression of exon 5 of *Crybb1* in eye tissue. In **C** and **D**, the x-axis shows genome position, and the-y axis denotes RNA-seq mean depth of samples in different genotype groups.

The outbred HS rat population has been extensively used in addiction-related behavioral studies, so we examined whether the eVariants catalog from our joint eQTL analyses contribute to the genetic architecture of addiction-related traits. Using published HS rats GWAS studies on heroin vulnerability^43^, cocaine self-administration^46^, cue-reactivity^44^, delay-discounting^42^ and open field behaviors^41^, we investigated the relationship between our joint eQTL results and 69 GWAS lead SNPs. Because the HS rats have extensive LD^48^, making colocalization extremely difficult, we instead asked whether the eQTL variants are tagged by published GWAS lead SNPs. Using LD estimated in our 86 outbred HS rats, we identified 80 lead eVariants in strong LD (r^2^ ≥ 0.8) and within ±1 Mb of 23 GWAS lead SNPs, yielding 111 GWAS-eQTL pairs (Table S6). Of these 80 lead eVariants, 42 were non-SNP variants (28 INDELs, 11 STRs, 3 SVs). As expected for cis-regulatory architectures, these lead eVariants were mostly noncoding.

Broadening to all eVariants at the same thresholds yielded 29,139 eVariants across 37 GWAS lead SNPs (206,094 GWAS-eQTL pairs), including 4,489 non-SNP eVariants (Table S6). A small subset showed exonic consequences (218 SNPs, 39 INDELs, 3 STRs, 1 SV), mostly in UTRs.

To illustrate the kinds of non-SNP variants implicated, we highlight two exonic eVariants that are strongly LD-tagged by GWAS lead SNPs. We focused on exonic variants because, unlike intronic variants, they provide a clear molecular mechanism for altering gene expression. First, we considered an 83-bp SV deletion in the 3’ UTR of *Zfp9* that is associated with decreased *Zfp9* expression in PL, IL and OFC tissues. This SV lies 67,031 bp from and is in strong LD (r^2^ = 0.8799) with the lead SNP for mean and total distance in the social zone (Figure S15A). Second, an AAAT-repeat STR in the 5’ UTR of *Get1* shows that longer repeat alleles are associated with increased *Get1* expression in PL, IL, and OFC tissues. It is 404,241 bp from and in strong LD (r^2^ = 0.9277) with the lead SNP for total heroin consumption (Figure S15B). Although the extensive LD precludes fine mapping, the fact that these variants are exonic makes them plausible explanations for these complex trait associations.

## Discussion

In this study, we used whole-genome short and long reads to generate the most comprehensive catalog of genetic variants to date for the HS rat population, integrating SNPs, indels, STRs, and SVs. By leveraging this multi-variant resource for joint cis-eQTL mapping across five addiction-relevant brain regions, we demonstrated that diverse variant types beyond SNPs made substantial contributions to regulatory variation. Our results showed that while SNPs remained the most abundant variant type overall, indels, STRs and SVs collectively accounted for more than 50% of the lead regulatory associations. Crucially, nearly 30% of these non-SNP lead eVariants were poorly tagged by nearby SNPs, underscoring that a meaningful fraction of associations is likely to be missed in SNP-only eQTL analysis due to incomplete genetic variant coverage. Direct comparison to SNP-only eQTL analyses showed that over 46% of shared eGenes were mapped to a non-SNP lead eVariant in the joint eQTL analyses. Moreover, over 53% of these variants were not in strong LD with their corresponding lead eSNPs in the SNP-only eQTL analyses. This highlights that a significant portion of regulatory signals detected in the SNP-only setting actually have stronger association evidence from non-SNP variants. Incorporating these complex variant types is therefore essential to identifying putative causal variants.

By including complex variant classes, we were able to discover regulatory mechanisms that were not identified when mapping eQTLs with only SNPs. A compelling demonstration is the 220 bp eSV deletion we identified in *Crybb1*, which disrupts both exonic and adjacent intronic sequence, causing dosage-dependent exon skipping. Our integration of published SNP-based GWAS with this new joint eQTL catalog revealed widespread overlap between addiction-related loci and eQTLs. By evaluating non-SNP eVariants strongly tagged by GWAS lead SNPs, we identified novel regulatory contexts for previously reported loci. The eSV deletion in *Zfp9* adds *Zfp9* as a novel candidate gene for social interaction behaviors that index anxiety and are predictors of vulnerability to addiction^41,72–74^. In humans, the ortholog *ZNF248* has been implicated in antidepressant class response and treatment-resistant depression^75^. The eSTR in *Get1* adds on the previously reported SNP-to-*Get1* splice-QTL to further strengthen *Get1* as a candidate gene for heroin consumption^43^. These examples illustrate complementary routes by which non-SNP variation influences complex traits, expanding the biological interpretation of existing GWAS signals.

Our variant discovery framework is consistent with recent advances in human and mouse genomics showing that long-read sequencing substantially improves the discovery of complex genetic variation beyond SNPs^51,76–79^. In humans, PacBio HiFi and other long-read approaches have enabled accurate detection of small variants, STRs and SVs, and pangenome studies have further shown that many complex or repetitive-region variants are incompletely represented by short-read sequencing against a single linear reference^51,76,78^. Similar long-read studies in mice have revealed extensive structural variation across diverse strains, including variants that affect local genome architecture and gene regulation^62,80^. In rats, recent long-read efforts have improved reference genome quality and variant discovery, but population-scale integration of SNPs, indels, STRs, and SVs has remained limited^81,82^. Thus, our combined short- and long-read variant catalog extends these advances to the HS rat population and provides a more complete representation of genetic variation for downstream functional mapping.

Our joint cis-eQTL analyses also build on studies in humans and other species showing that non-SNP variants contribute meaningfully to gene expression variation^5,16,17,83–85^. Previous work has demonstrated that SVs and STRs can act as regulatory variants and that joint analyses incorporating multiple variant classes can identify associations not well captured by SNP-only approaches^5,16,65,84,85^. Consistent with these findings, we observed that indels, STRs, and SVs collectively accounted for a substantial fraction of lead eVariants across five addiction-relevant brain regions, and many of these signals were poorly tagged by nearby SNPs. These results indicate that SNP-only eQTL analyses can miss or misprioritize regulatory variants when the underlying causal signal is driven by more complex forms of genetic variation. By applying a unified multi-variant eQTL framework in HS rats, our study provides evidence that incorporating indels, STRs, and SVs improves the resolution of regulatory mapping in a genetically diverse model population.

While this study establishes a foundational resource, several limitations should be considered when interpreting the results. First, the relatively low PacBio HiFi coverage (∼10x) for the 86 HS rats restricted our focus to common variants that passed stringent filtering. We could not comprehensively catalog rare variants or fully explore de novo variant discovery. Second, relying on a linear reference genome (mRatBN7.2) likely caused us to miss or misrepresent variants in highly complex genomic regions enriched for SVs and STRs. Large intragenic STRs and SVs can also interfere with standard RNA-seq read mapping using a linear reference genome, meaning our measured expression levels may not perfectly capture true transcript abundance for some severely disrupted genes. Adopting a pangenome reference in future studies would better capture this structural diversity and improve mapping accuracy in these complex and polymorphic regions^78,86–91^. Third, encoding bi-allelic SNPs, indels, and SVs as 0/1/2 dosages and STRs as normalized allele length sums was necessary for joint analysis, but may have introduced biases in effect size estimation or reduced sensitivity to true regulatory effects. Fourth, extensive LD blocks in the HS rat population limited our ability to perform fine-mapping and colocalization. Because many STRs and SVs are poorly tagged by SNPs, our ability to definitively connect specific eQTLs with GWAS signals remains constrained.

In summary, we generated a comprehensive multi-type genetic variant and eVariant catalog for the HS rat population, providing a foundational resource to study the role of different types of variants in complex traits. Future studies that impute these variants into HS rats for which only SNP genotypes derived from low pass short read sequencing are available will enable genetic mapping studies that use all variant types. We anticipate that such studies would identify new associations and may also improve associations previously attributed to SNPs by demonstrating that other variant classes showed stronger associations. Ultimately, our results offer clear evidence that SNP-only eQTL mapping provides an incomplete understanding of genetically determined differences in gene regulation; this conclusion is not limited to HS rats but extends to other model organisms.

## Methods

### Subjects

The N/NIH HS rat is an outbred population originally developed in 1984 by interbreeding eight inbred “founder” strains (ACI/N, BN/SsN, BUF/N, F344/N, M520/N, MR/N, WKY/N, and WN/N)^35,39^. Since its inception the outbred HS rats population has been maintained by interbreeding a large number of families per generation for over 100 generations, allowing the founder haplotypes to be progressively broken down by recombination and providing a well-powered framework for genetic mapping ^36–39^.

Two cohorts of rats were used in this study. For each cohort, we used preexisting short read sequencing data and generated new long read sequencing data. The first cohort consisted of tissues that had been continuously frozen since 1984 from the 8 inbred founder strains that were used to create the HS populations. The second cohort consisted of 86 outbred HS rats from generations 73-80 that were the focus of a previous eQTL mapping study^48^.

### SNPs and indels

The bi-allelic SNPs and indels for the eight founders and 86 outbred HS rats used in this study were generated from Illumina whole-genome sequencing (**WGS**) short reads and were previously published^49^. The mean sequencing coverage for the eight founder strains (8 males) is 41.81x (NCBI SRA: PRJNA1048943), and 33.24x for the 86 outbred HS rats (44 females and 42 males; NCBI SRA: PRJNA1076141) (Figure S1A). To avoid indels’ overlaps with other types of variants in this study, we filtered out indels with greater than 50% reciprocal overlap with STRs, and indels with length greater than 50 bp (Figure S2). 9,710,761 bi-allelic SNPs and 1,893,961 bi-allelic indels were retained after filtering using BCFTools^92^.

### Long reads sequencing

Whereas the short read data used in this paper had been generated previously, we generated new long read sequence data for this paper. We performed Pacific Biosciences (**PacBio**) high fidelity (**HiFi**) long-read sequencing. DNA was extracted using PacBio Nanobind PanDNA kits (previously Circulomics Nanobind Big DNA kits). For the founder strains, we extracted DNA from small intestine (BN/SsN, BUF/N, M520/N), liver (ACI/N, MR/N, WKY/N, WN/N) or tail (F344/N) tissues. For the 86 outbred HS rats, their spleen tissues were used to extract DNA. DNA libraries were prepared using PacBio HiFi plex prep kits according to the manufacturer’s protocol. The DNA was then sequenced with PacBio Sequel IIe platform at the University of California San Diego Institute for Genomic Medicine Genomics Center or PacBio Revio at the Stem Cell Genomics Core at the Sanford Stem Cell Institute. The mean sequencing coverage for the eight founder strains (1 female and 7 males) is 41.05x (NCBI SRA: PRJNA1048943), and 10.51x for the 86 outbred HS rats (44 females and 42 males; NCBI SRA: PRJNA1076141) (Figure S1A).

Due to the poor tissue quality of the historical eight founders, F344/N and MR/N sequenced with PacBio HiFi were not the exact samples as the ones with Illumina WGS. To be specific, two different male F344/N samples were sequenced. For MR/N, a male sample was used for Illumina WGS and a female sample for PacBio HiFi. For the 86 outbred HS rats, the exact same individuals were used for Illumina WGS, PacBio HiFi, as well as bulk RNAseq for generating the gene expression dataset mentioned in the Methods section.

The PacBio HiFi long reads were mapped to Rattus norvegicus reference genome mRatBN7.2 (NCBI Genome Assembly Accession: GCF_015227675.2) using minimap2^93,94^ (minimum mapping quality: 20, minimum alignment length: 1 kb). Next, before the variant calling process, the aligned long reads were used to phase SNPs generated from short reads using Whatshap^95^, and the phased SNPs were used to tag the haplotypes in long reads (Figure S3A and S4A).

### Short tandem repeats

The STRs were jointly genotyped for the eight founders and 86 outbred HS rats using aligned PacBio HiFi long reads (Figure S3A). We, first, used Tandem Repeats Finder^96^ (match=2, mismatch=7, delta=7, PM=80, PI=10, minscore=5, maxperiod=500, -l 6) to identify the set of reference STRs with repeat unit length between 1-6bp from the Rattus norvegicus reference genome mRatBN7.2. Segmental duplication regions were identified using Parascopy^97^ and removed from the reference STRs set. LongTR^98^ was used to genotype STRs (--min-reads 5, -- max-tr-len 10 kb).

After obtaining the initial STR callset, we applied a set of call-level filters to ensure genotypes’ quality (Figure S3B and S3C). Pentanucleotides with homopolymer runs of 5 bp or longer and hexanucleotides with homopolymer runs of 6 bp or longer were filtered out because these long homopolymer tracts are prone to sequencing and alignment errors, leading to excess indels and unreliable STR genotype calls. Due to the different PacBio HiFi long reads coverage between HS founders (41.05x) and outbred HS rats (10.51x), different filters were used on coverage related filters. For HS founders, only genotypes with call coverage between 10 and 100 and minimum four supporting reads for each allele were kept. For outbred HS rats, only genotypes with call coverage between 4 and 100 and minimum one supporting reads for each allele were retained. We then filtered out genotypes with quality score less than 0.9, homopolymer and dinucleotide STRs that are heterozygous in more than 80% of the genotyped inbred founders, and STRs with no length variation across all samples. This resulted in 724,032 STRs. TRTools^99^ was used for STR filtering.

### Structural variants

Joint SV calling was performed with the eight founders and 86 outbred HS rats using aligned PacBio HiFi long reads (Figure S4A). Sniffles2^100^ was used to conduct single-sample SV calling followed by a multi-sample joint SV calling. During the single-sample calling process, variants with supporting reads less than 0.1 x (0.25 x average chromosomal coverage + 0.75 x variant surrounding coverage) were discarded. Variant calls with fewer than three minimum variant reads support were also discarded. Sniffles2 detects each allele separately and separates multi-allelic SVs into different bi-allelic SVs. To support our downstream eQTL analysis, we kept the obtained SVs as bi-allelic.

A set of call-level filters were also applied to the initial SV callset to remove low-confidence calls (Figure S4B and S4C). Breakend (BND) SVs, SVs with length less than 50 bp or greater than 1 Mb, and SVs with greater than 50% reciprocal overlap with STRs were filtered out. We further filtered out SV variant calls with genotype quality less than five, resulting in 115, 371 SVs for the HS rat population. BCFTools^92^ was used for SV filtering.

### Variants quality control

After the various variant calling and call-level filters mentioned above, we also filtered out variants based on locus-level metrics. We separately filtered out variants with missing rate > 0.1 for the eight HS founders and the 86 outbred HS rats. Because the 86 outbred HS rats were used for eQTL mapping, we further filtered out their variants to ensure the power for the association tests. First, we examined if any variant violated Hardy-Weinberg Equilibrium (**HWE**). For SNPs, indels and SVs, who are biallelic, the ones with -log10(HWE *P*-value) > 10 were filtered out. For STRs, which tend to be highly multiallelic, we used their observed and expected percent of homozygous calls to compute HWE *P*-value, and filtered out the ones with - log10(HWE *P*-value) > 5. Although the outbred HS rats used in this study were from the same population, and a lenient HWE threshold would be sufficient, we applied a more stringent threshold for STRs because they tend to have higher genotyping error rate than other types of variants. The HWE filter was only applied to autosomal variants for all samples and chromosome X variants for female samples. Next, we only kept common variants from the 86 outbred HS rats. SNPs, indels and SVs with minor allele frequency less than 0.05 were filtered out. For STRs, we filtered out variants with heterozygosity less than 0.1. The heterozygosity is positively correlated with non-major allele frequency in STRs, and STRs with non-major allele frequency less than 0.05 all had heterozygosity less than 0.1, so an additional allele frequency filter for STRs was not needed. Plink2^101^, BCFTools^92^ and TRTools^99^ were used for these variant filtering steps. Detailed variant filtering processes can be found in Figure S2B, S2C, S3B, S3C, S4B and S4C. As a result, we retained 8,744,914 bi-allelic SNPs, 1,649,180 bi-allelic indels, 167,884 STRs and 94,778 SVs for eight HS founders, and 5,944,671 bi-allelic SNPs, 1,025,924 bi-allelic indels, 134,917 STRs and 46,550 SVs for 86 outbred HS rats.

### STRs evaluation

To evaluate the obtained STRs, we compared the obtained STR callset after filtering to another set of STRs produced by Illumina WGS short reads using a short-read calling pipeline with HipSTR^102^ (Figure S5A, S5B and S5C). To be specific, genotype concordance was calculated by comparing the length difference between genotyped and reference alleles (GB field produced by LongTR and HipSTR) while allowing one base-pair difference. We computed the genotype concordance rate separately for the eight HS founders and 86 outbred HS rats. For the eight HS founders, for which we have 41.05x PacBio HiFi reads, the number of shared STRs between long-read calling and short-read calling pipelines was 85,256 with an average concordance rate of 96.95% (Figure S5D and S5E). For the 86 outbred HS rats, for which we only have 10.51x PacBio HiFi reads, the number of shared TRs between long-read calling and short-read calling pipelines was 81,076 with an average concordance rate of 92.53% (Figure S5F and S5G).

### SVs evaluation

To evaluate the obtained SVs using an orthogonal data modality approach, we utilized the available 10.8x coverage PacBio HiFi and 41.1x coverage Oxford Nanopore sequences of one outbred HS rat to construct an SV truth set, and compared this sample’s SV genotypes produced by the joint SV calling pipeline to the truth set (Figure S6A). Specifically, two sets of orthogonal long-read data were separately passed through the same SV calling pipeline with the same set of call-level filters mentioned above, generating two independent sets of SVs. The SV truth set for this outbred HS rat was then built by extracting SVs with greater than 50% reciprocal overlap between these two sets of SVs. Because the truth set was produced by merging calls from two single-sample calling pipelines, in order to do an equal comparison, the SVs before locus-level filtering (missing rate, HWE, minor allele frequency filters) from the SV joint calling pipeline were extracted for comparison. Compared to the truth set, there were 70,909 SVs detected by the joint calling pipeline that were not in the truth set (less than 50% reciprocal overlap); however, the majority of them (60,449 SVs, 85.2%) had homozygous reference genotypes, which is a result of joint calling. Of the 38,115 SVs in the truth set, 88.2% (33,632 SVs) were captured by the SV joint calling pipeline (equal to or greater than 50% reciprocal overlap) with a genotype concordance rate of 87.68% (Figure S6B and S6C).

### Gene expression dataset

The gene expression data of the 86 outbred HS rats used in this study were curated from RatGTEx portal (https://RatGTEx.org v3 unmerged version). Detailed data preprocessing procedures were described on the RatGTEx portal and our previous publication^48^. In this study, we used inverse-quantile normalized gene expression data from five brain tissues: prelimbic cortex (**PL**, n=78, 40 females and 38 males), infralimbic cortex (**IL**, n=79, 40 females and 39 males), orbitofrontal cortex (**OFC**, n=78, 39 females and 39 males), nucleus accumbens core (**NAcc**, n=73, 38 females and 35 males) and lateral habenula (**LHb**, n=80, 40 females and 40 males). The data was mapped to Rattus norvegicus reference genome mRatBN7.2 (NCBI Genome Assembly Accession: GCF_015227675.2) and annotated with RefSeq gene annotations. We also further filtered out genes not expressed in more than half of the samples. As a result, we obtained 19,389 genes for PL, 19,315 genes for IL, 19,275 genes for OFC, 19,109 for NAcc and 19,451 genes for LHb.

### eQTL mapping

Because of the differences among the different types of variants, genotypes were pre-processed before entering the eQTL analysis. In this study, we used bi-allelic SNPs and indels; hence, their genotypes were encoded to 0, 1, 2 as dosage for the eQTL analysis input. Since the SV calls outputted by the Sniffles2 caller are also bi-allelic, SV genotypes were processed in the same way as SNPs and indels. For STRs, which tend to be more multi-allelic, we used the normalized sum of allele lengths as dosage input, which is scaled to be between 0 and 2. Because different tissues used in this study had different sample sizes, we further filtered out variants with greater than 0.1 missing rate and less than 0.05 non-major allele frequency for each tissue before the eQTL mapping analysis.

We adapted the eQTL mapping pipeline used for the RatGTEx^48^ and performed cis-eQTL mapping using tensorQTL^103,104^ on genotype dosages described above. Gene expression levels were tested for association with variants located within a ±1 Mb window around each gene’s transcription start site. The first 20 principal components of the gene expression phenotypes and first 10 principal components of the pruned genotypes were included as covariates. Empirical *P*-values were estimated using beta approximation from permutation derived null distributions for each gene. These were used to compute gene level q-values, applying a false discovery rate threshold of 0.05 to identify genes with significant associations (**eGenes**). For each significant gene, nominal *P*-value thresholds derived from the permutations were used to determine significant variants (**eVariants**).

## Resource availability

### Lead contact

Requests for further information and resources should be directed to and will be fulfilled by the lead contact, Abraham Palmer.

### Materials availability

This study does not generate new unique reagents.

HS rats are available at https://ratgenes.org/cores/core-b/.

### Data and code availability

Eight HS inbred founder Illumina short and PacBio long reads are available at NCBI SRA: PRJNA1048943.

Eighty-six outbred HS rat Illumina short and PacBio long reads are available at NCBI SRA: PRJNA1076141.

Outbred HS rat RNAseq raw reads are available at NCBI SRA: PRJNA723867, and expression data are available at RatGTEx portal (https://ratgtex.org/, v3 unmerged).

Genotype data after quality control and eQTL mapping summary stats are available at UC San Diego Library Digital Collections (https://doi.org/10.6075/J05X29WF).

Variant calling, eQTL mapping and analysis code is available at GitHub (https://github.com/Palmer-Lab-UCSD/HS-Rats-Complex-Variants-eQTL-Paper).

## Acknowledgments

This work was supported by the National Institutes of Health grant P30DA060810 (A.A.P.), U01DA051234 (M.G., J.L.S. and A.A.P.), P50DA037844 (A.A.P.) and F31DA063333 (D.C.). PacBio Sequel IIe HiFi long-reads sequencing was conducted at the IGM Genomics Center, University of California, San Diego, La Jolla, CA. We would like to thank the Stem Cell Genomics Core at the Sanford Stem Cell Institute for providing PacBio Revio HiFi long-reads sequencing services. We would also like to thank Oxford Nanopore Technologies for providing one outbred HS rat ONT long reads sequencing through a pilot project. San Diego Supercomputer Center (2022): Triton Shared Computing Cluster. University of California, San Diego. Service. https://doi.org/10.57873/T34W2R

## Author contributions

Conceptualization, D.C., M.G., J.L.S., A.A.P.

Data Curation, D.C., K.A.C., K.H.N., Y.W., T.M., M.M., D.M., O.P.;

Formal Analysis, D.C., H.Z.J., R.J.E.

Funding Acquisition, D.C., M.G., J.L.S., A.A.P.

Methodology, D.C., H.Z.J., J.G., M.M., D.M.

Supervision, O.P., M.G., J.L.S., A.A.P.

Visualization, D.C.

Writing – Original Draft, D.C.

Writing – Review & Editing, All authors

## Declaration of generative AI and AI-assisted technologies in the writing process

During the preparation of this work, the authors used ChatGPT and Gemini to help revise the original text for readability. After using this tool, the authors reviewed and edited the content as needed and take full responsibility for the content of the publication.

## Supplemental information titles

Document S1: Figures S1-S15

Document S2: Tables S1-S6

