## Supplementary Figures S1-S15 for "Point mutations and complex variants impact gene expression and addiction-related behaviors in Heterogeneous Stock rats"

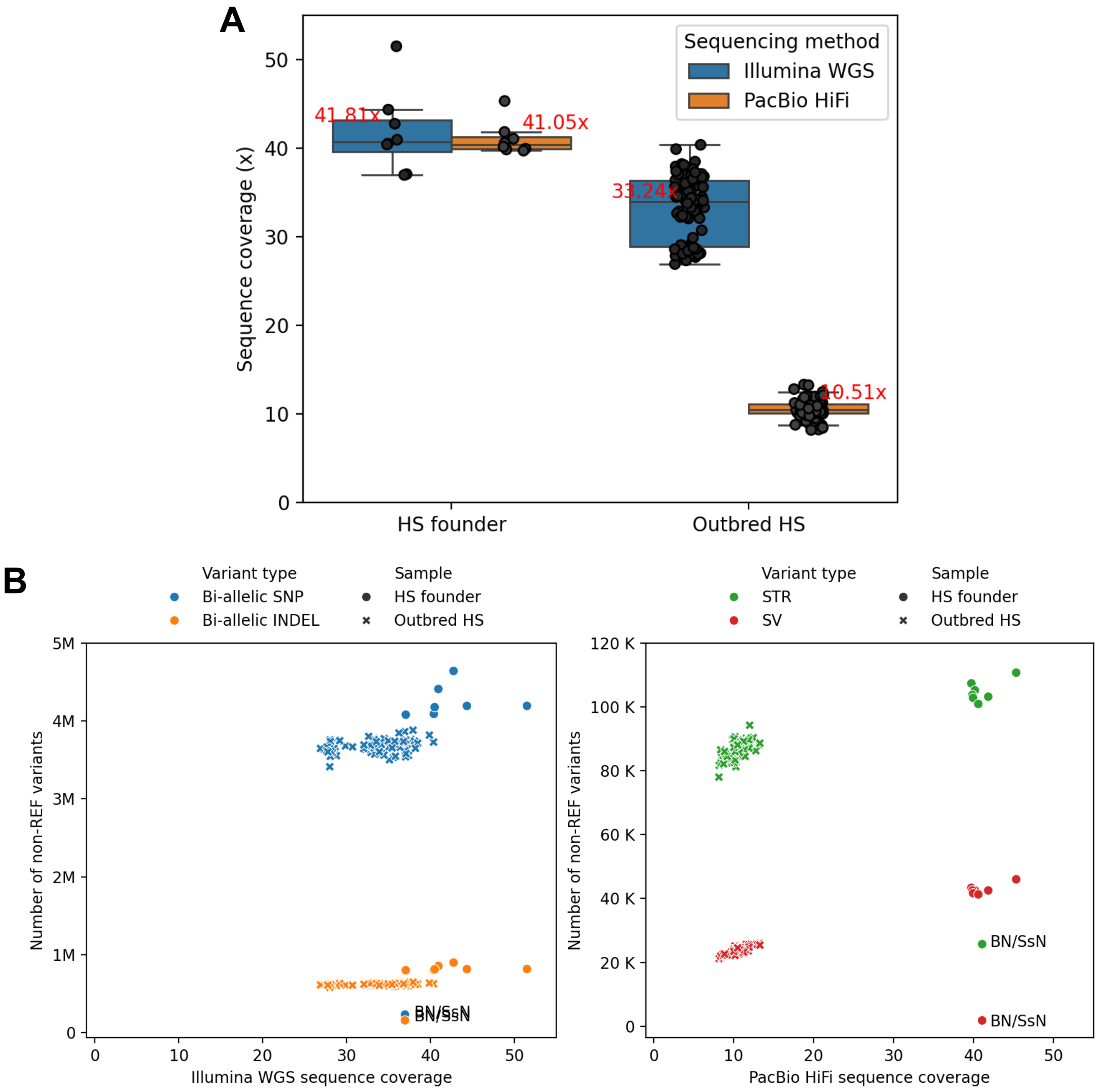


**Supplementary Figure S1**: **A.** Sample DNA sequence coverage. Each dot is a sample. Annotated red text denotes the mean of each box. **B.** Number of variants detected for each sample. Each dot is a sample. BN/SsN, a reference genome like strain, is annotated.

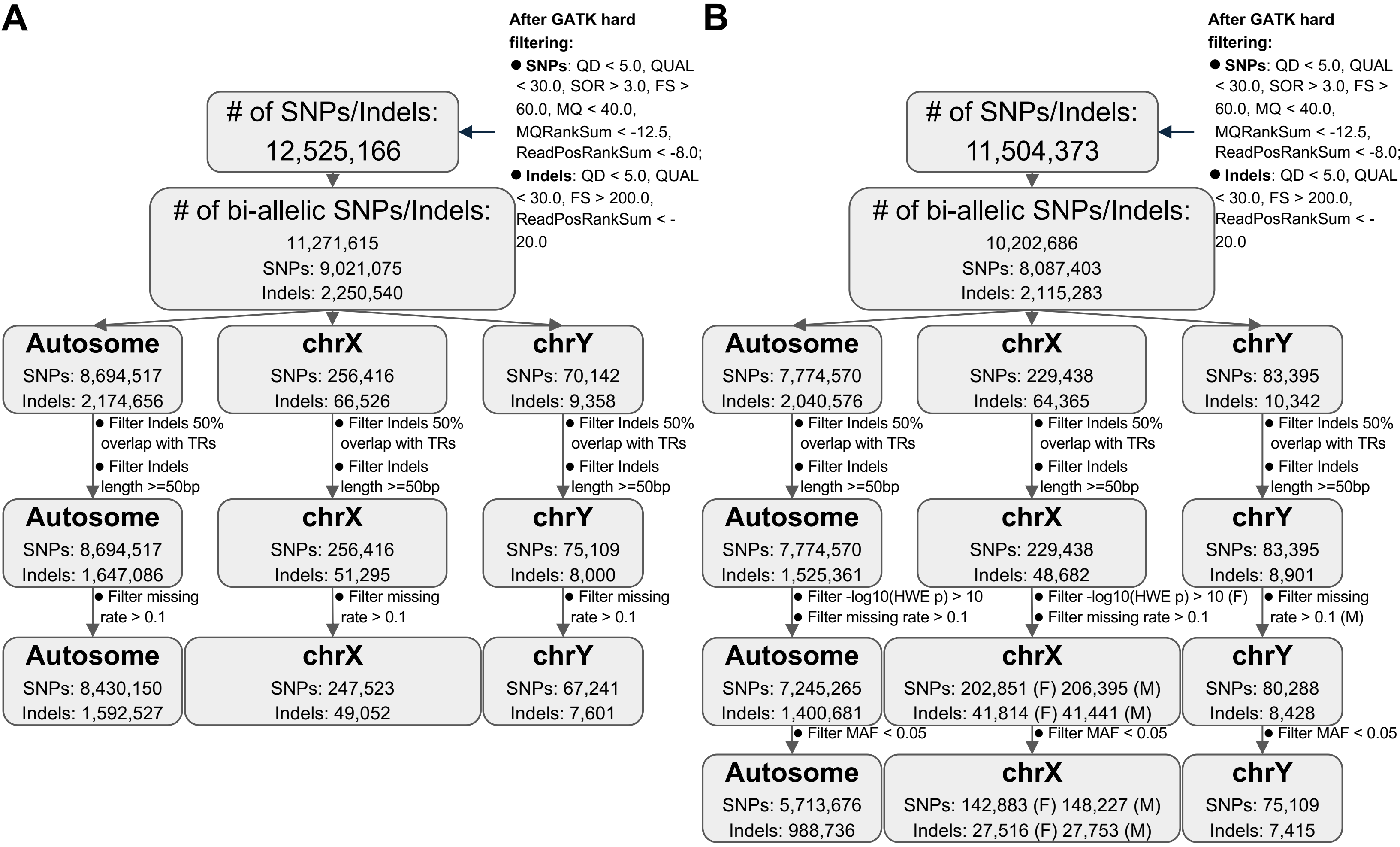


**Supplementary Figure S2**: SNPs and indels quality control. **A.** Eight HS founders. **B.** 86 outbred HS rats.

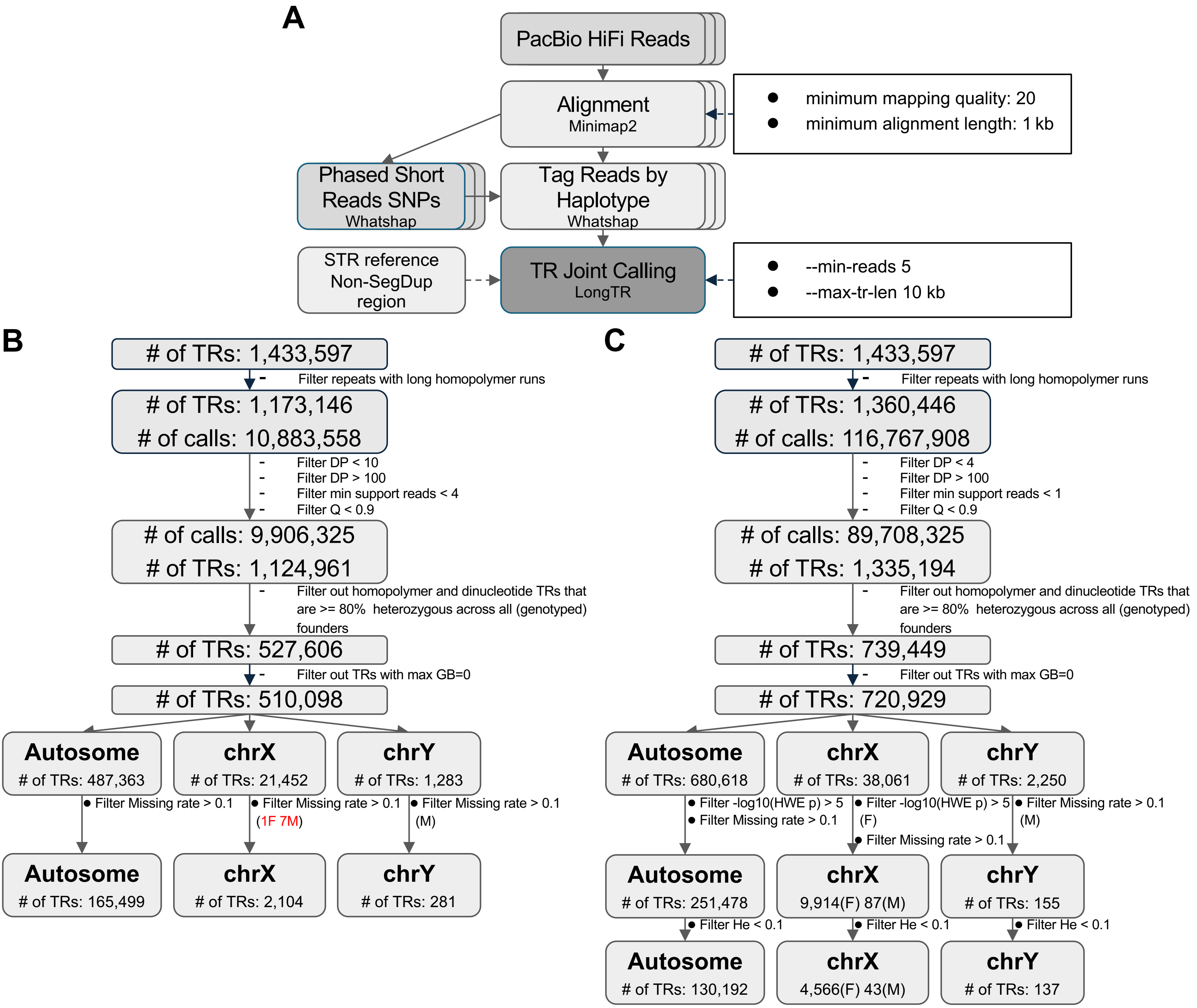


**Supplementary Figure S3**: STR calling pipeline and quality control. **A.** STR calling pipeline flowchart. **B.** STR call-level and locus-level quality control for eight HS founders. **C.** STR call-level and locus-level quality control for 86 outbred HS rats.

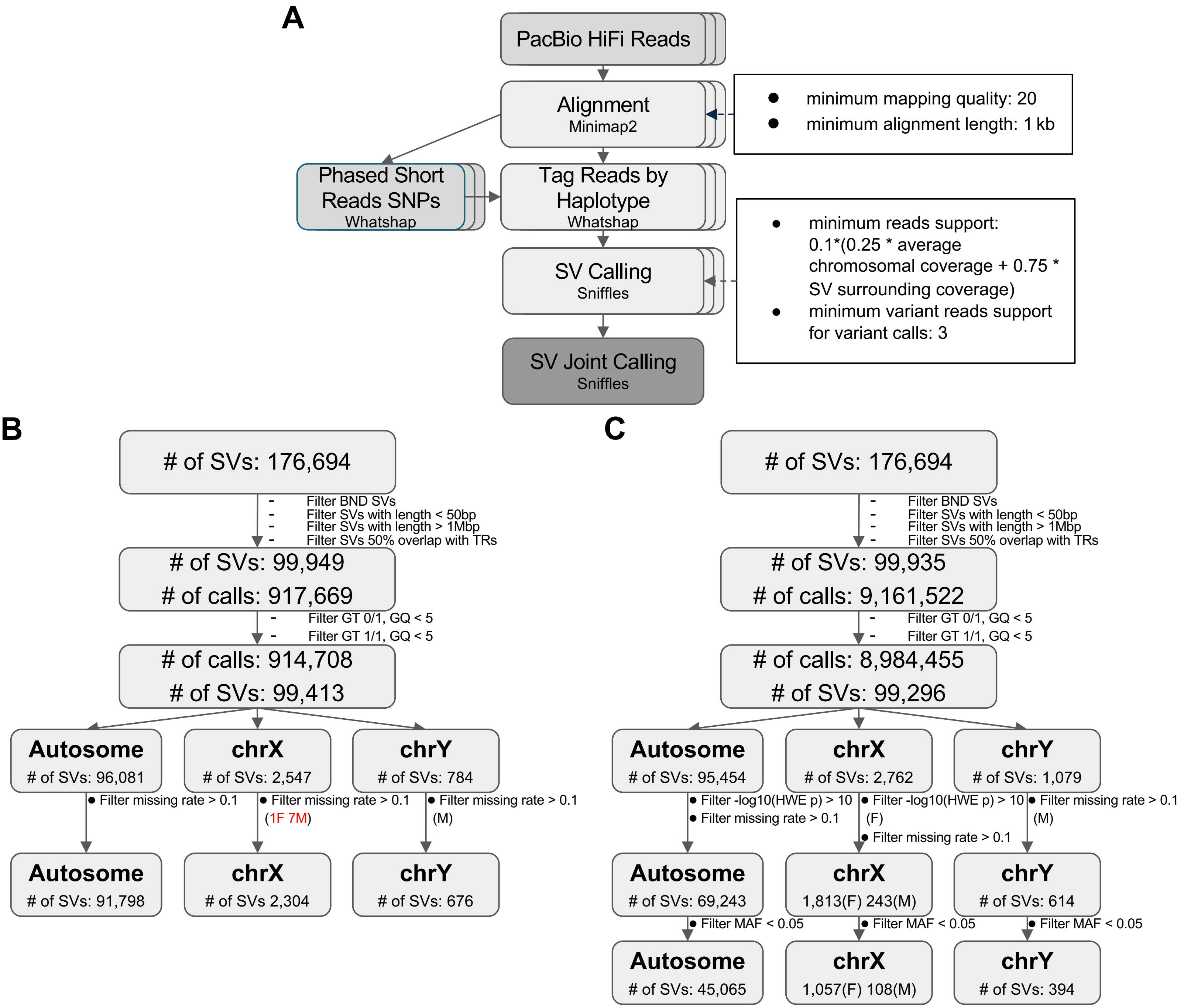


**Supplementary Figure S4**: SV calling pipeline and quality control. **A.** SV calling pipeline flowchart. **B.** SV call-level and locus-level quality control for eight HS founders. **B.** SV call-level and locus-level quality control for 86 outbred HS rats.

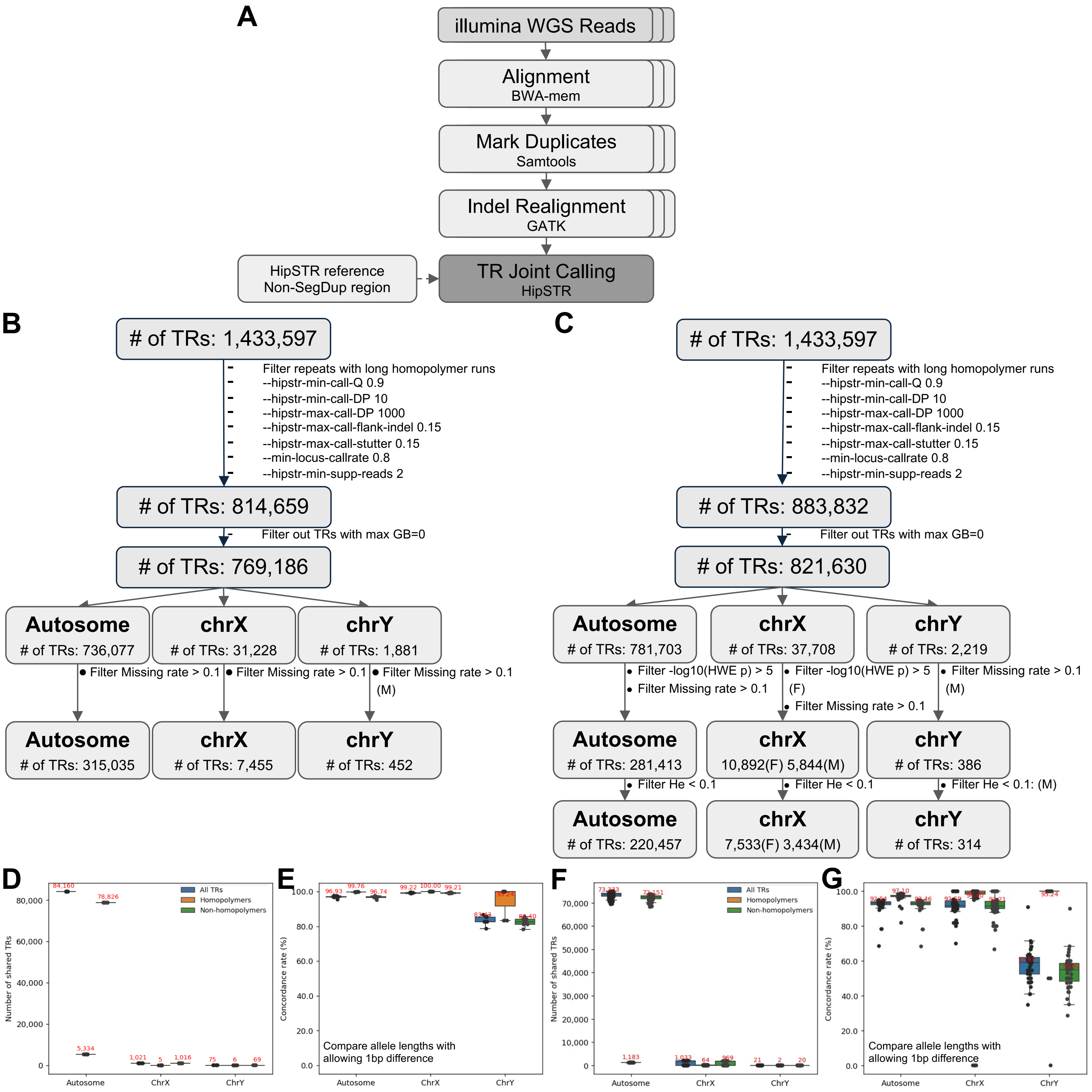


**Supplementary Figure S5**: STR evaluation. **A.** STR short-read calling pipeline flowchart. **B.** STR short-read calling pipeline quality control for eight HS founders. **C.** STR short-read calling pipeline quality control for 86 outbred HS rats. **D.** Number of shared STRs between long-read calling and short-read calling pipeline for eight HS founders. **E.** Concordance rate of the genotyped STRs' GB fields between long-read calling and short-read calling pipeline for eight HS founders. **F.** Number of shared STRs for 86 outbred HS rats. **G.** Concordance rate of the genotyped STRs' GB fields for 86 outbred HS rats. In **D/E/F/G**, each dot is a sample, and annotated red text denotes the mean of each box.

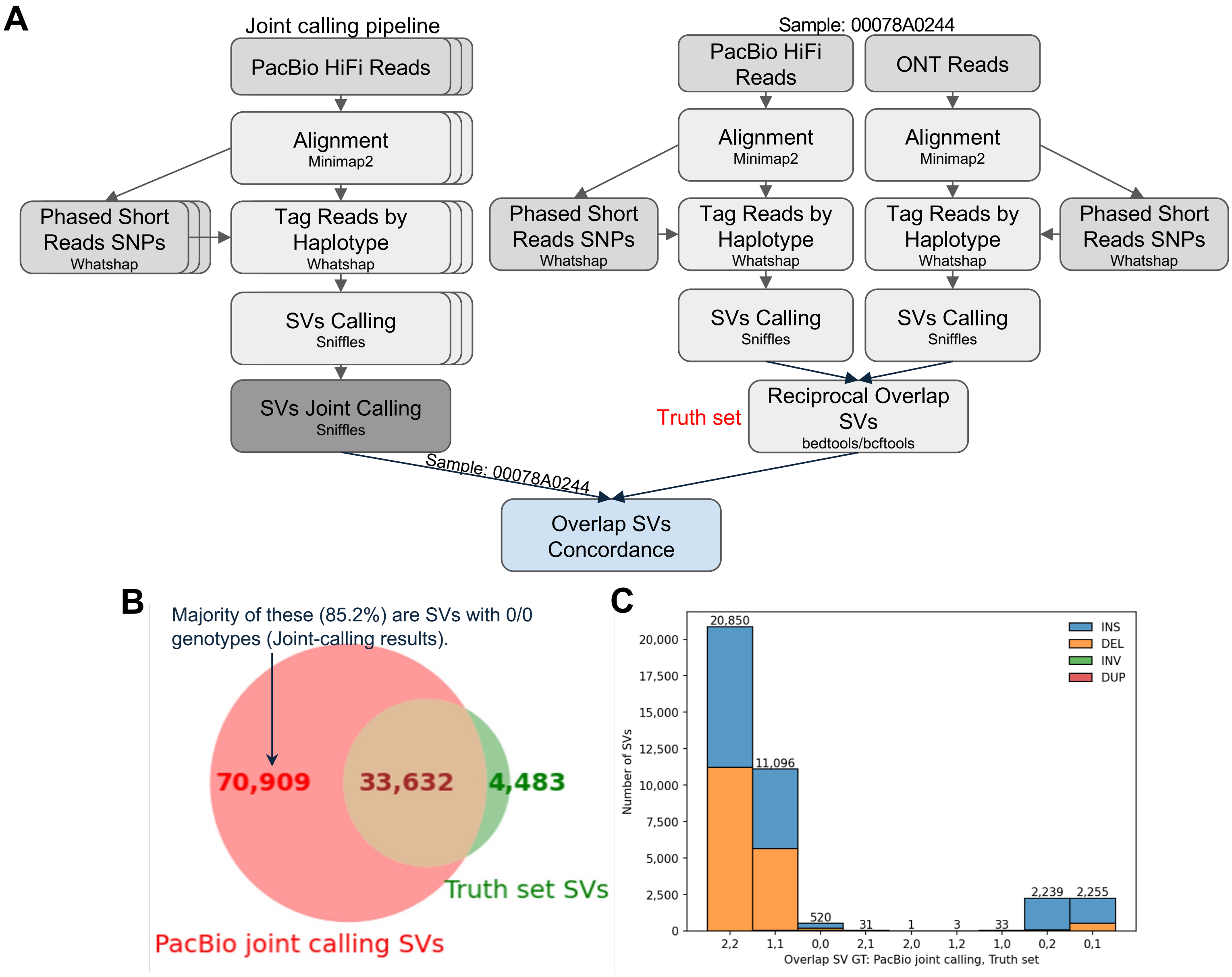


**Supplementary Figure S6**: SV evaluation. **A.** SV evaluation strategy. **B.** Number of shared SVs between joint SV calling pipeline and the truth set. **C.** Genotype concordance check for shared SVs between joint SV calling pipeline and the truth set.

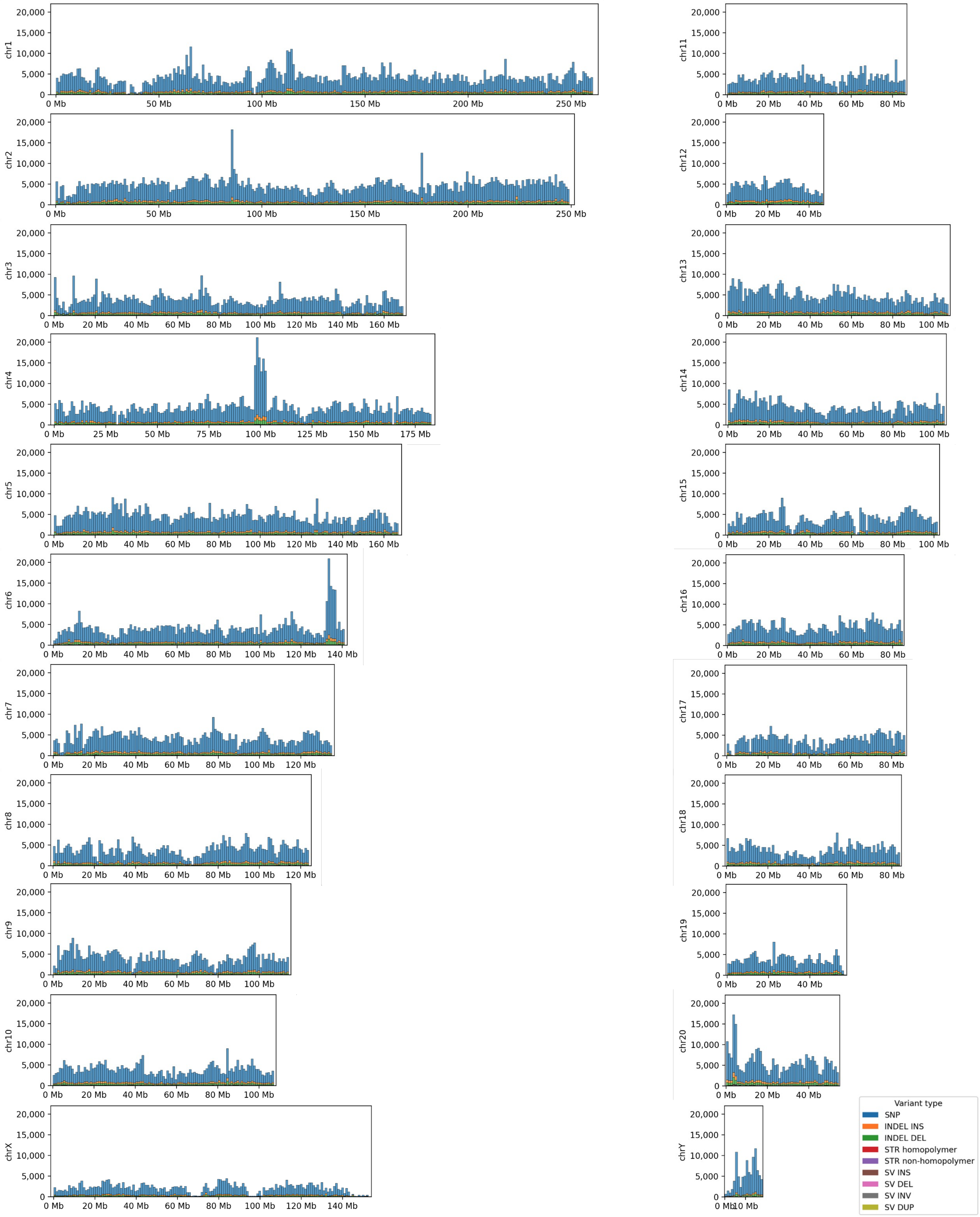


**Supplementary Figure S7**: Variant density distribution histogram on each chromosome with one-megabase windows.

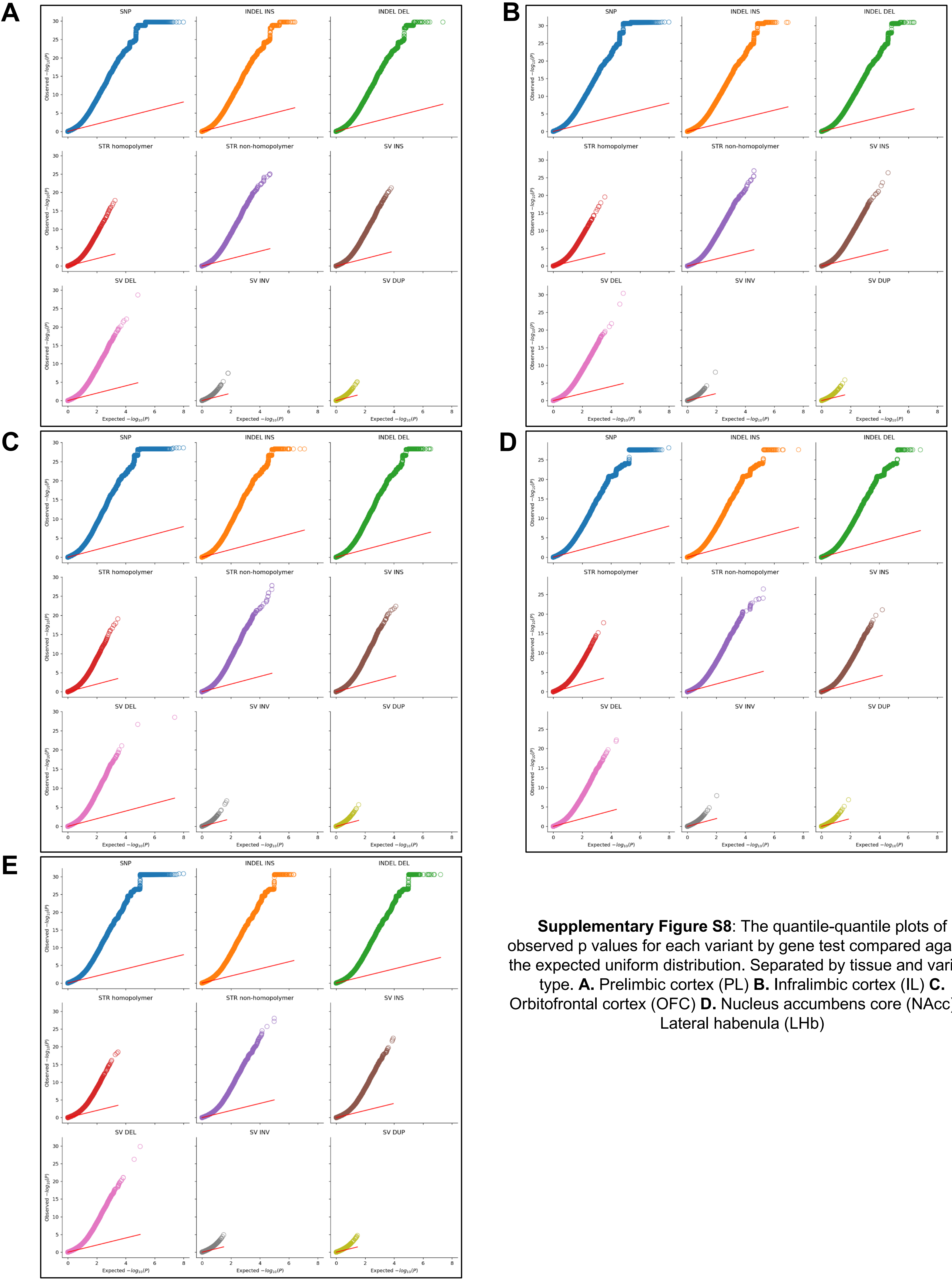


**Supplementary Figure S8**: The quantile-quantile plots of observed p values for each variant by gene test compared against the expected uniform distribution. Separated by tissue and variant type. **A.** Prelimbic cortex (PL) **B.** Infralimbic cortex (IL) **C.** Orbitofrontal cortex (OFC) **D.** Nucleus accumbens core (NAcc) **E.** Lateral habenula (LHb)

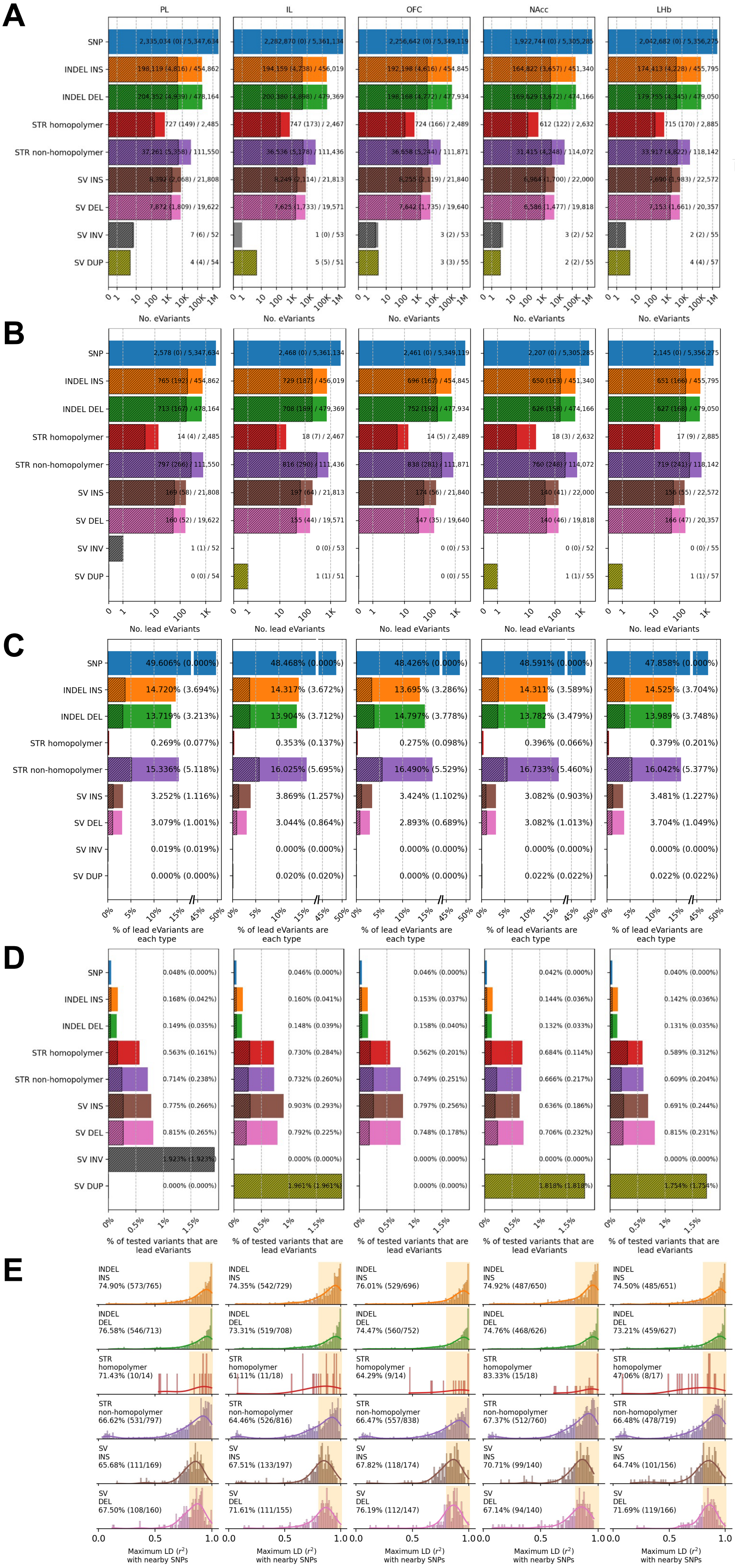


**Supplementary Figure S9**: eVariants statistics across different tissues. **A.** Number of eVariants detected. **B.** Number of lead eVariants. **C.** Percent of lead eVariants that are each variant type. **D.** Percent of tested variants from each variant type that are lead eVariants. **E.** Maximum linkage disequilibrium distribution between a non-SNP lead eVariant and SNPs within one megabase.

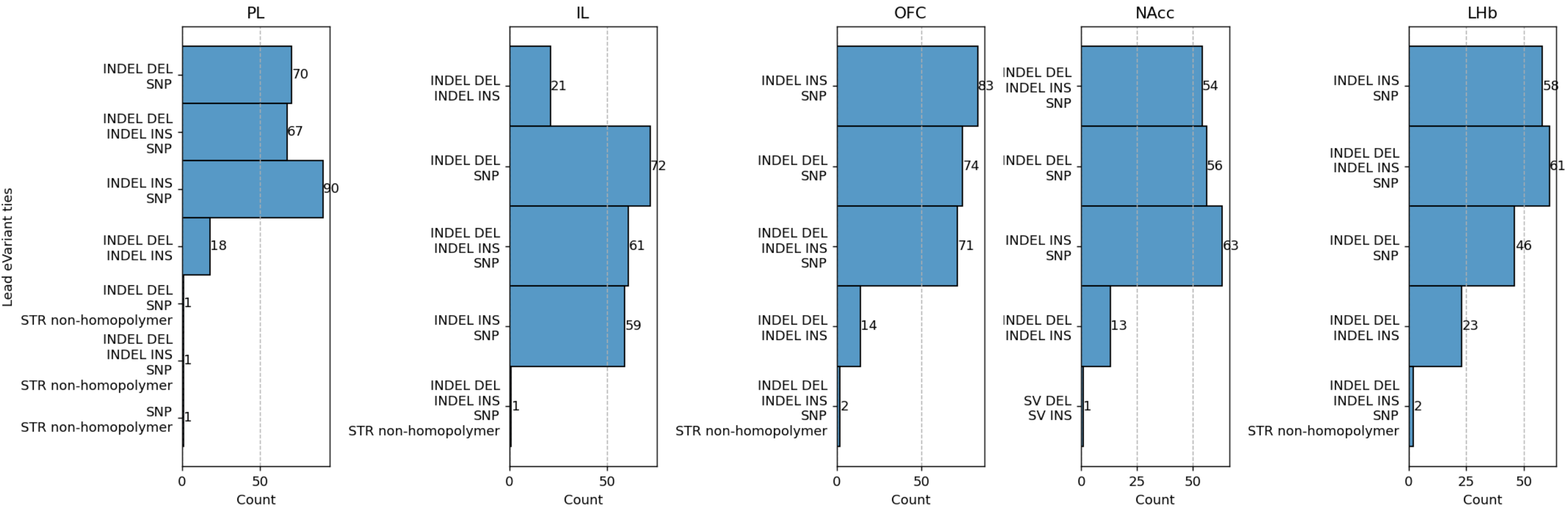


**Supplementary Figure S10**: Number of lead eVariants ties in each tissue.

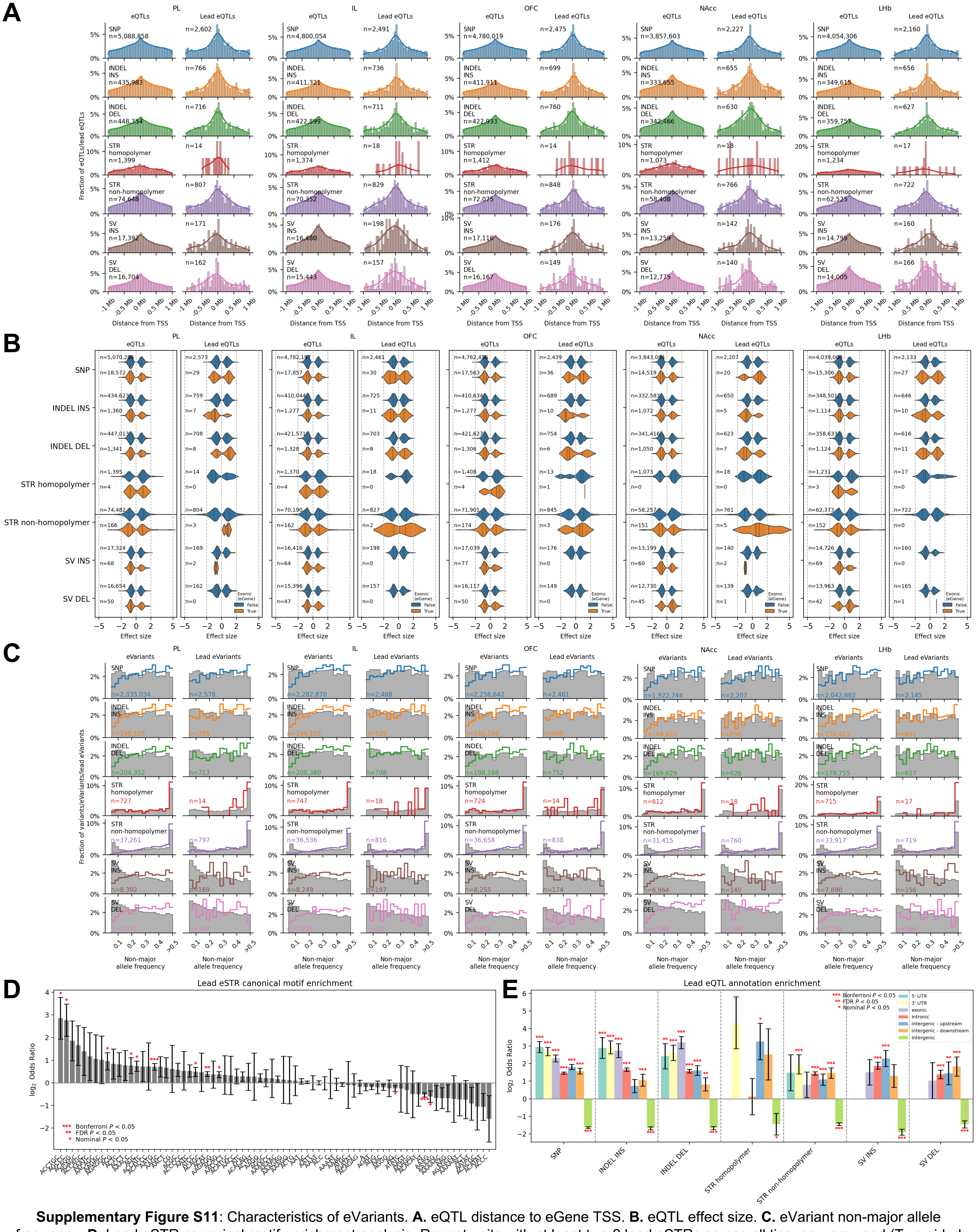


**Supplementary Figure S11**: Characteristics of eVariants. **A.** eQTL distance to eGene TSS. **B.** eQTL effect size. **C.** eVariant non-major allele frequency. **D.** Lead eSTR canonical motif enrichment analysis. Repeat units with at least two 2 lead eSTRs across all tissues were used (Two-sided Fisher's exact test against all tested STRs. Error bars represent ±1 s.e.m.). **E.** Lead eQTL annotation enrichment analysis (Two-sided Fisher's exact test against all tested variants. Error bars represent ±1 s.e.m.).

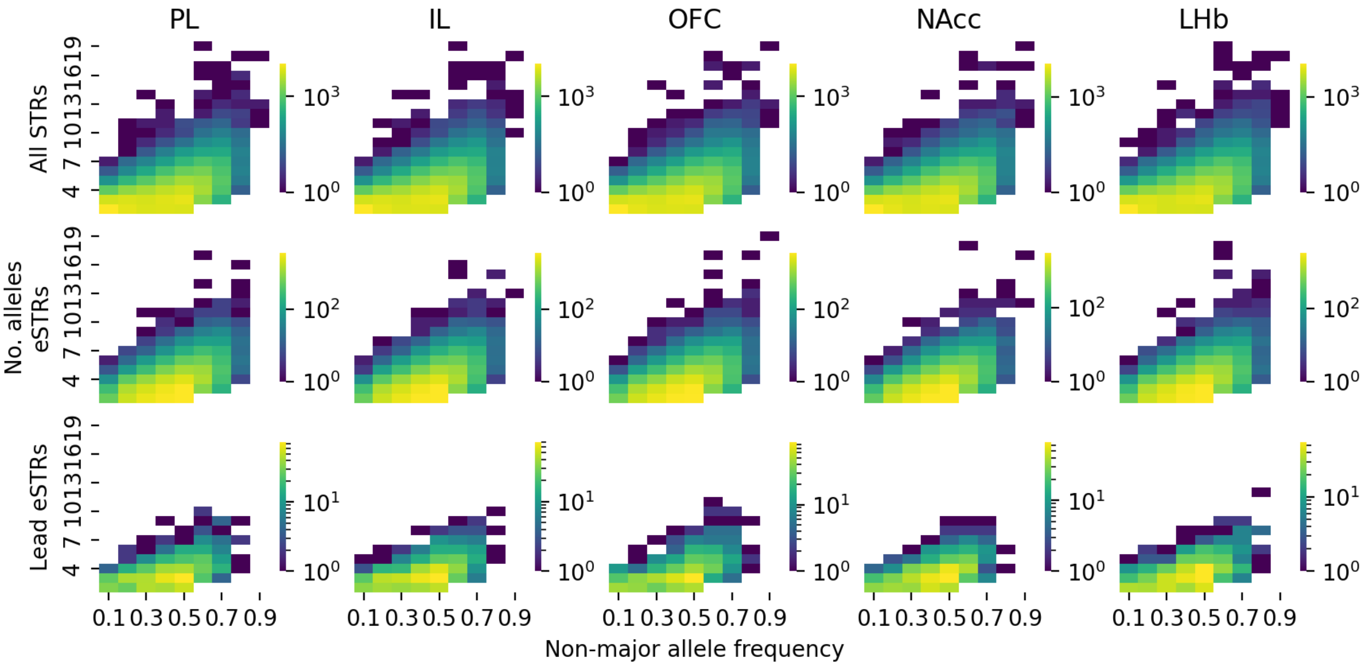


**Supplementary Figure S12**: STR number of alleles vs. non-major allele frequency.

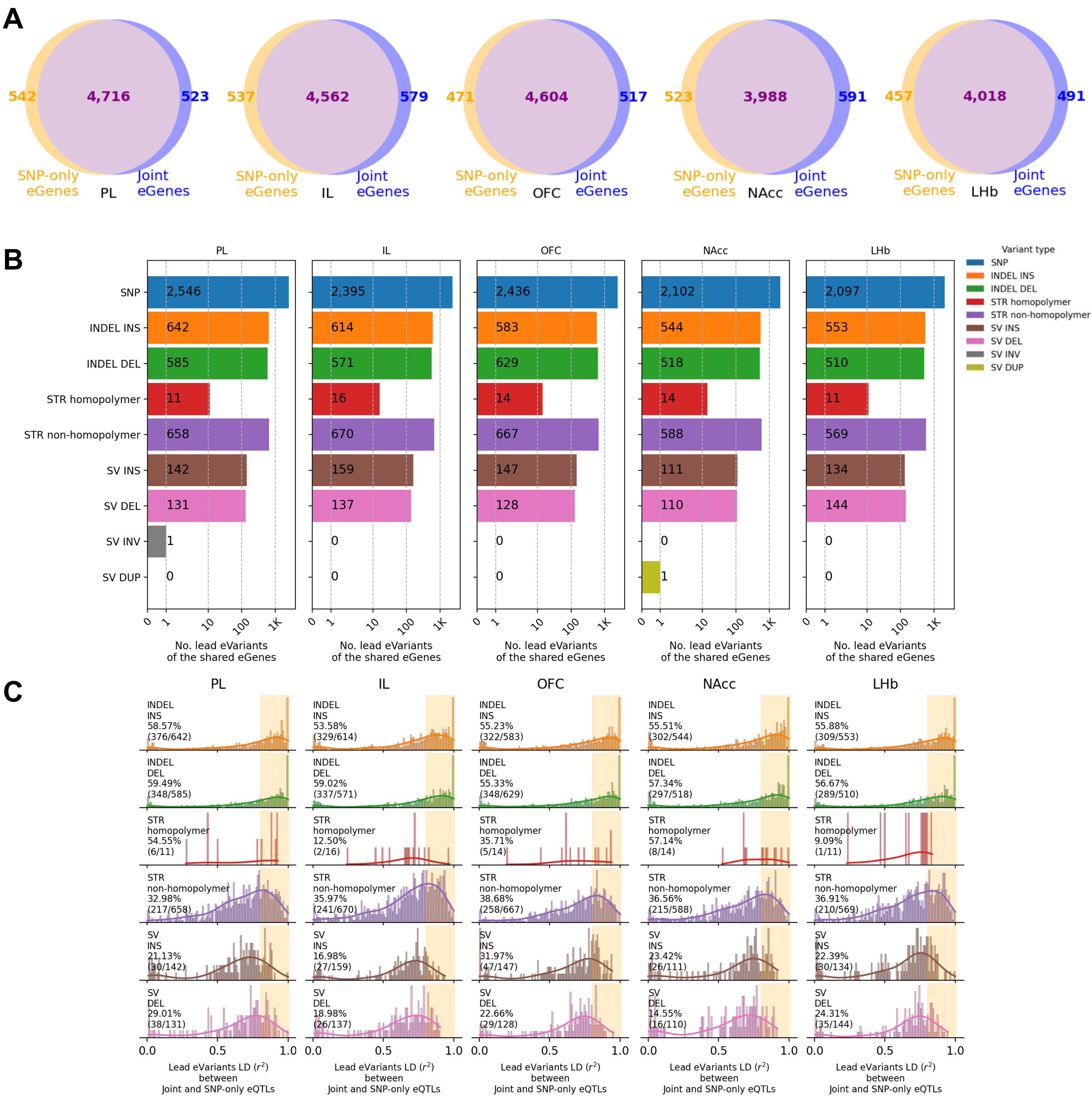


**Supplementary Figure S13**: Joint versus SNP-only eQTL analyses across different tissues. **A.** Venn diagram showing the intersection between eGenes detected in the joint and the SNP-only eQTL analyses. **B.** Number of lead eVariants of the shared eGenes across variant types. **C.** Linkage disequilibrium distribution between shared eGenes' lead eVariants in the joint eQTL analysis and corresponding lead eSNPs in the SNP-only eQTL analysis.

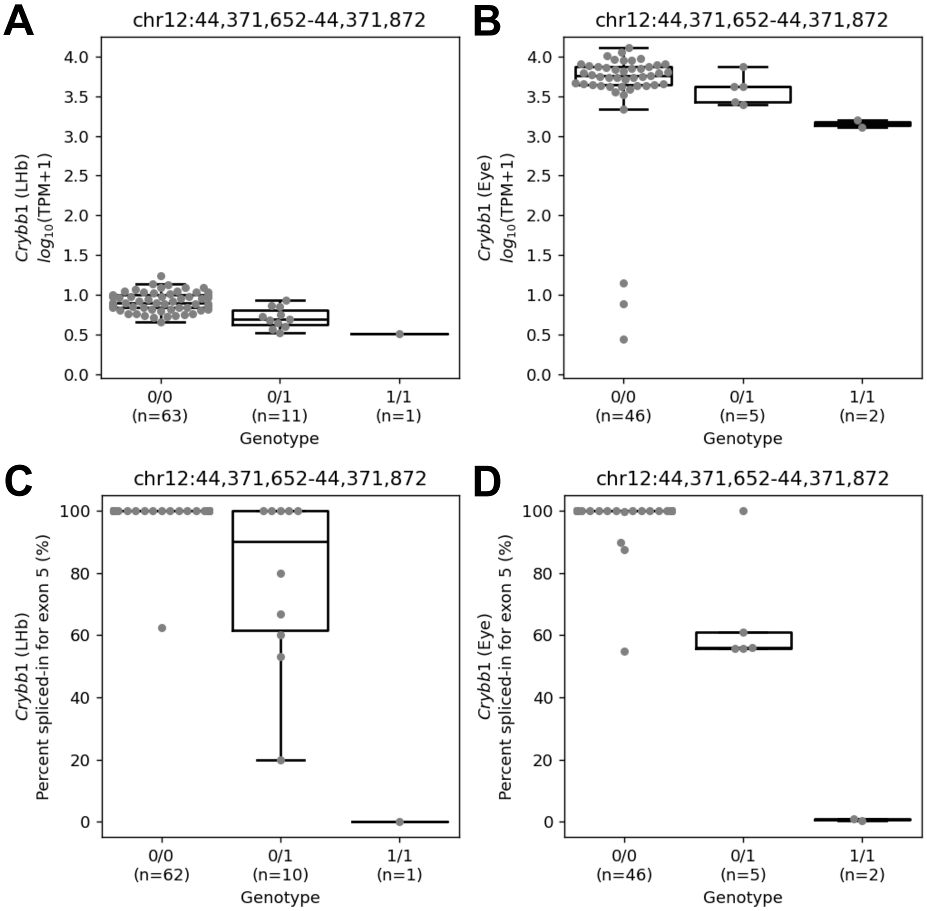


**Supplementary Figure S14**: Exon-disrupting eSV affecting expression of *Crybb1* in LHb and eye tissues. **A.** Effect plot for the SV-eQTL in LHb tissue. **B.** Effect plot for the SV-eQTL in eye tissue. In **A** and **B**, the x-axis shows genotype groups with sample sizes, and the y-axis shows -$\log_{10}$(TPM+1). **C**. Percent spliced-in for *Crybb1* exon 5 in LHb tissue. **D.** Percent spliced-in for *Crybb1* exon 5 in eye tissue. In **C** and **D**, samples with zero total reads from exon 4-5, exon 5-6 and exon 4-6 junctions were filtered out.

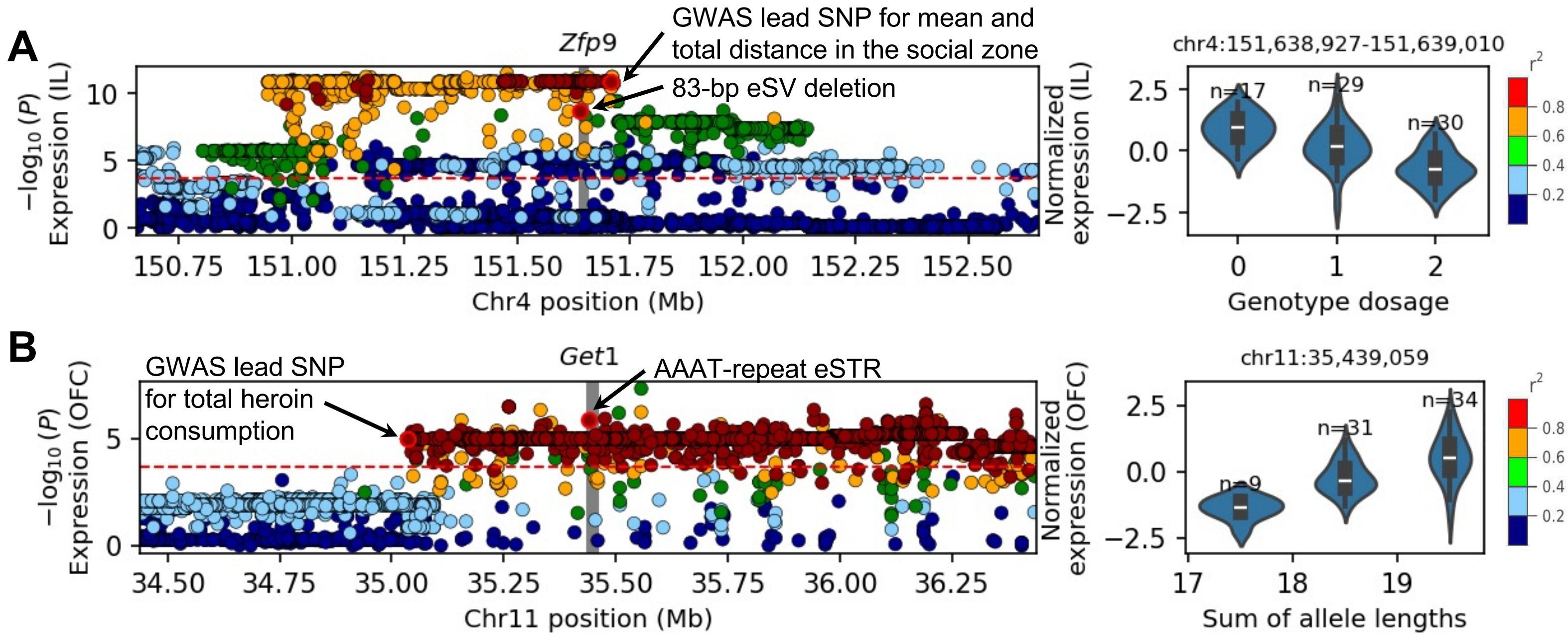


**Supplementary Figure S15**: Candidate non-SNP eQTLs at GWAS loci. **A.** An 83-bp eSV deletion in 3' UTR of Zfp9 was in strong LD with HS rat GWAS lead SNP for mean and total distance in the social zone. **B.** An AAAT-repeat eSTR in 5' UTR of Get1 was in strong LD with HS rat GWAS lead SNP for total heroin consumption. In **A** and **B**, the left panels show the nominal -$\log_{10}$($P$-values) for all tested variants for the corresponding gene. The gray background indicates the genomic position of the gene. Point colors represent the LD $r^2$ between each variant and the GWAS lead SNP. The right panels show the corresponding eQTL effect plot for with genotypes on the x-axis and normalized expression on the y-axis.
